# A novel isoform of *ACE2* is expressed in human nasal and bronchial respiratory epithelia and is upregulated in response to RNA respiratory virus infection

**DOI:** 10.1101/2020.07.31.230870

**Authors:** Cornelia Blume, Claire L Jackson, Cosma Mirella Spalluto, Jelmer Legebeke, Liliya Nazlamova, Franco Conforti, Jeanne-Marie Perotin-Collard, Martin Frank, Max Crispin, Janice Coles, James Thompson, Robert A Ridley, Lareb S N Dean, Matthew Loxham, Adnan Azim, Kamran Tariq, David Johnston, Paul J Skipp, Ratko Djukanovic, Diana Baralle, Chris McCormick, Donna E Davies, Jane S Lucas, Gabrielle Wheway, Vito Mennella

## Abstract

Angiotensin-converting enzyme 2 (ACE2) is the main entry point in the airways for SARS-CoV-2. ACE2 binding to SARS-CoV-2 protein Spike triggers viral fusion with the cell membrane, resulting in viral RNA genome delivery into the host. Despite ACE2’s critical role in SARS-CoV-2 infection, an understanding of ACE2 expression, including in response to viral infection, remains unclear.

Until now *ACE2* was thought to encode five transcripts and one 805 amino acid protein. Here we identify a novel short isoform of ACE2. Short *ACE2* is expressed in the airway epithelium, the main site of SARS-CoV-2 infection; it is substantially upregulated in response to interferon stimulation and RV infection, but not in response to SARS-CoV-2 infection, and it shows differential regulation in asthma patients. This short isoform lacks SARS-CoV-2 spike glycoprotein high-affinity binding sites and altogether, our data are consistent with a model where short *ACE2* may influence host susceptibility to SARS-CoV-2 infection.

## Introduction

At the time of writing there have been more than 15 million confirmed cases of COVID-19 and more than 600,000 confirmed COVID-19 associated deaths worldwide (WHO; 26^th^ July 2020). There is therefore an urgent global need to understand the molecular mechanism of infection and disease to identify patients’ susceptibility and targets for therapeutic intervention.

A key molecule responsible for SARS-CoV-2 viral entry is the metalloprotease angiotensin-converting enzyme 2 (ACE2), a transmembrane protein encoded by the human *ACE2* gene. *ACE2* consists of 19 exons and encodes five annotated transcripts, two of which encode the same 805 amino acid protein ACE2 (UniProt Q9BYF1) which has a theoretical predicted molecular mass of 92.4 kDa and observed mass of ~120 kDa due to multiple sites of glycosylation of the N-terminus region^1^. ACE2 consists of an N-terminal extracellular domain and a C-terminal membrane anchor domain^1^. The extracellular domain contains a 17-amino acid signal peptide sequence, an N-terminal catalytic metallopeptidase domain with 41% amino acid sequence identity to ACE^1,2^ and a C-terminal domain with 48% amino acid sequence identity to renal amino acid transporter collectrin (TMEM27)^3^. The extracellular domain can be shed via cleavage of specific residues in the ferredoxin-like fold domain (neck dimerization domain) of the protein by ADAM17, TMPRSS11D or TMPRSS2 proteases. TMPRSS2 expression and activity has been shown to increase SARS-CoV-2 viral entry^4,5^.

ACE2 is a carboxypeptidase with several known physiological functions. It catalyses the removal of the C-terminal residue from different vasoactive peptides, its key substrate being angiotensin II, which contribute to the renin-angiotensin system, a physiological feedback loop regulating blood pressure, salt and water balance in mammals^1,2^. In the small intestine ACE2 is co-expressed and interacts with amino acid transporter B(0)AT1 (SLC6A19) at the brush border to form a catalytic complex required for amino acid uptake^6,7^. By homology to collectrin, ACE2 is also required for glucose homeostasis and pancreatic beta-cell function^8,9^.

Interestingly, ACE2 plays an important role in protection from acute lung injury. *Ace2* expression is downregulated in mice models of acute lung injury and *Ace2* knockout mice show a more severe acute lung injury phenotype^10^. Moreover, improved outcomes are seen in pig models of lung injury in which *ACE2* is overexpressed^11^ and an activator of *ACE2*, XNT, can protect against pulmonary hypertension in rat models^12,13^. Although the molecular mechanism by which ACE2 protects against acute lung injury remains unclear, it is known that the carboxypeptidase function of ACE2 is required to confer this protection and that AT2 (AngII receptor 2) also confers protection^10^.

Importantly, ACE2 is the main viral entry point for coronavirus N63, SARS-CoV and SARS-CoV-2, which cause severe-acute respiratory syndromes, the latter being responsible for COVID-19 in humans^14–17^. ACE2 binds to the S1 domain of trimeric SARS-CoV-2 Spike glycoprotein^18^, and viral entry is dependent upon the extracellular domain of ACE2 being cleaved by TMPRSS2 protease at arg697 and lys716^19^, and the transmembrane domain of ACE2 internalised with the virus via the clathrin-mediated^20,21^ and clathrin-independent^22^ endocytosis pathways.

*ACE2* expression in different tissues is controlled by multiple promoter elements^23^. *ACE2* expression is under the control of Ikaros homology activating elements around −516/−481 in the heart^24^ and under the control of estrogen responsive elements in adipose tissue^25^. *ACE2* expression in human nasal epithelia and lung tissue is under the control of interferon-responsive promoters, with STAT1, STAT3, IRF8, and IRF1 binding sites at −1500–500 bp*^26^*. Activation of interferon (IFN) responsive genes is an important antiviral defence pathway in humans, and both interferon and influenza exposure lead to an increase in *ACE2* expression in human airway^26^ most likely reflecting its anti-inflammatory role which serves to protect against acute lung injury following viral infection.

Bulk RNA sequencing data^27^ detects low-level expression of *ACE2* in testis, small intestine, thyroid, colon, kidney, heart left ventricle and atrial appendage, and visceral adipose. Single cell RNA sequencing (scRNAseq) studies show *ACE2* expression at low levels in airway, cornea, esophagus, ileum, colon, liver, gallbladder, heart, kidney and testis^28^. Using scRNAseq and RNA *in situ* hybridisation, *ACE2* expression in the airways has been observed to be relatively high in nasal epithelium and progressively lower in the bronchial and alveolar regions^29^. Highest expression is seen in goblet and ciliated cells of the nasal epithelium^28^, and ACE2 protein localises to the membrane of motile cilia of respiratory tract epithelia^30^. It has been demonstrated that SARS-CoV infection occurs through ACE2 on cilia in airway epithelia^30^. Airway multiciliated cells appear to be one of the main targets of SARS-CoV-2 infection^29^, suggesting that cilia are also important in SARS-CoV-2 infection, possibly because their extension from the cell surface makes ACE2 more accessible to the virus. Importantly, *ACE2* expression correlates with levels of infection of SARS-CoV-2 isolates from patients in different airway compartments^29^. These studies have established the upper airway as the main site of SARS-CoV-2 infection.

Here we detail the identification of a novel isoform of ACE2, which we name short ACE2, that is expressed in human nasal and bronchial respiratory epithelia, the main site of SARS-CoV-2 infection, and is preferentially expressed in asthmatic bronchial epithelium relative to full length ACE2 (long ACE2). In airway primary cells, short *ACE2* is upregulated in response to IFN-beta treatment and RNA respiratory rhinovirus infection, but not SARS-CoV-2.

## Results

### Identification of novel short ACE2 transcript in nasal and bronchial airway cells

We analysed the expression of *ACE2* in airway epithelia in our existing RNAseq datasets from nasal brushings and nasal epithelia cultured at air-liquid interface (ALI) by aligning reads to human genome build 38 using STAR 2-pass mapping^31^ and GENCODE v33 gene annotations. We visually analysed mappings to *ACE2* using Integrative Genomics Viewer^32^, which identified multiple reads mapping to a genomic region between exon 9 and 10 of the constitutive *ACE2* gene build (**Figure 1a**). These mappings showed a discrete 3’ junction at GRCh38 chrX: 15580281, but variable 5’ length suggesting a splice junction with downstream exon 10, but no splicing upstream to exon 8. This suggests that a novel unannotated exon exists between exon 9 and 10, and that this exon is the beginning of a novel transcript distinct from full-length *ACE2* transcripts ACE2-202 (ENST00000427411.1) or ACE2-201 (ENST00000252519.8) in the airway. Visual assessment of total read mappings to *ACE2* gene show approximately double the number of read support to exons 10-19 compared to exons 1-9, further suggesting that a novel shorter transcript of *ACE2*, which includes a novel exon plus exons 10-19, is expressed at equal or higher levels than longer ACE2 transcripts including exons 1-19 (ACE2-202) or 2-19 (ACE2-201) (**Figure 1a**). Assembly of transcriptomes from all samples using SCALLOP tool identified novel transcripts including this novel exon to exon 19 (**Figure 1b**). Sashimi plot analysis confirmed splicing between this new exon and downstream exon 10, but complete absence of splicing at the 5’ end of the new exon (**Figure 1c**). Analysis of these RNAseq data to identify and filter novel splice junctions with code developed by Cummings *et al*.^33^ also independently detected a novel splice junction at chrX: 15580281. Analysis of splice junctions identified by STAR aligner confirmed multiple uniquely-mapped reads to a novel exon/exon boundary removing a novel intron of coordinates GRCh38 chrX: 15578316–15580280. We call this novel exon 9a. Study of the sequence of the exon 9a/intron boundary showed a strong U1-dependent consensus splice site sequence (AG|GTAAGTA) suggesting that it is a strong splice donor site (**Figure 1d**). This splicing event introduces a new in-frame ATG start codon 29 nucleotides upstream of the splice site (**Figure 1d**), and a TATA box 148 nucleotides upstream of the splice site (**Figure 1d**) suggesting that this transcript is protein-coding. Furthermore, a promoter flanking region has been identified at GRCh38 chrX:15581200-15579724 (ENSR00000902026), suggesting active transcription upstream of exon 9a (approx. chrX:15580402 – chrX:15580281). Analysis of this region shows a near consensus ISGF-3 binding site (TgGTTTCAgTTTCCt)^34^ 159bp upstream of the splice junction, a near-consensus AP-1 binding site (TGtGTCA)^35^ 223bp upstream of the splice site and an NF-kB binding site (GGGTTTTCCC)^36^ 787bp upstream of the splice junction. This suggests that the short form of ACE2 is under independent transcriptional control from full-length ACE2 expression, and that this may be controlled by IFN, AP-1 and NF-kB elements.

**Fig 1.**
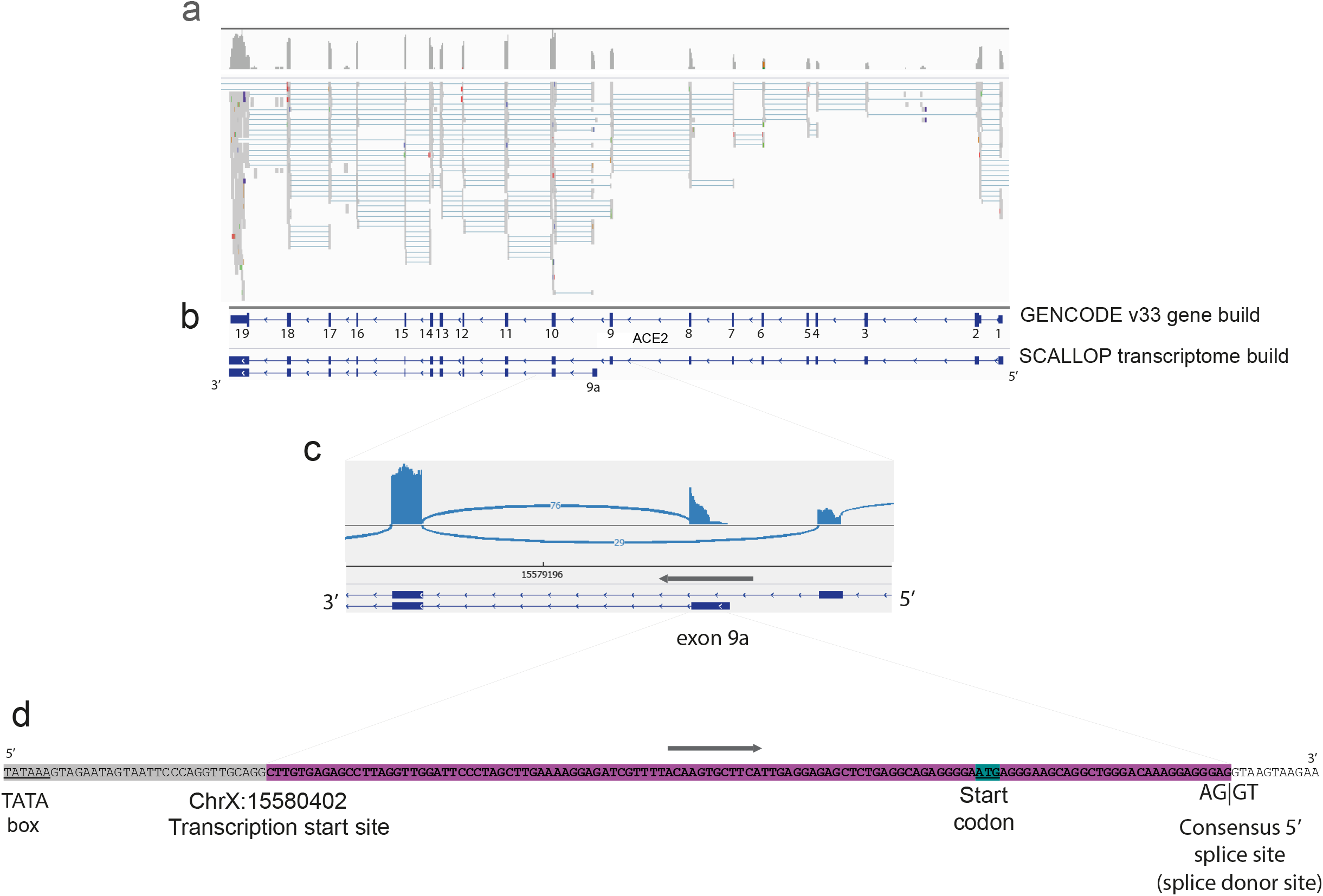
A novel short transcript of *ACE2* is expressed in airway epithelia. 1a. IGV plot showing reads mapped along ACE2 from one nasal brushing sample, including reads mapped to a region between exons 9 and 10. 1b. GENCODE v33 gene build exons and novel SCALLOP transcriptome build exons, showing novel exon 9a. 1c. Sashimi plot showing splice junction between exon 9a and 10 1d. Nucleotide sequence of novel exon 9a, plus 5’ UTR, start codon and splice junction

To confirm expression of this novel transcript we performed RT-PCR using primers specific to exon 1, exon 9a and exon 19 using cDNA from both nasal brushings and differentiated immortalized bronchial epithelial cells BCi-NS1.1^37^ (**Figure 2a, b**). We performed Sanger sequencing to confirm the identity of these PCR amplicons and confirmed sequences spanning exon 9 and 10, and exon 9a and 10 in the amplicons from the constitutive transcript and novel transcript, respectively (**Figure 2c**).

**Figure 2.**
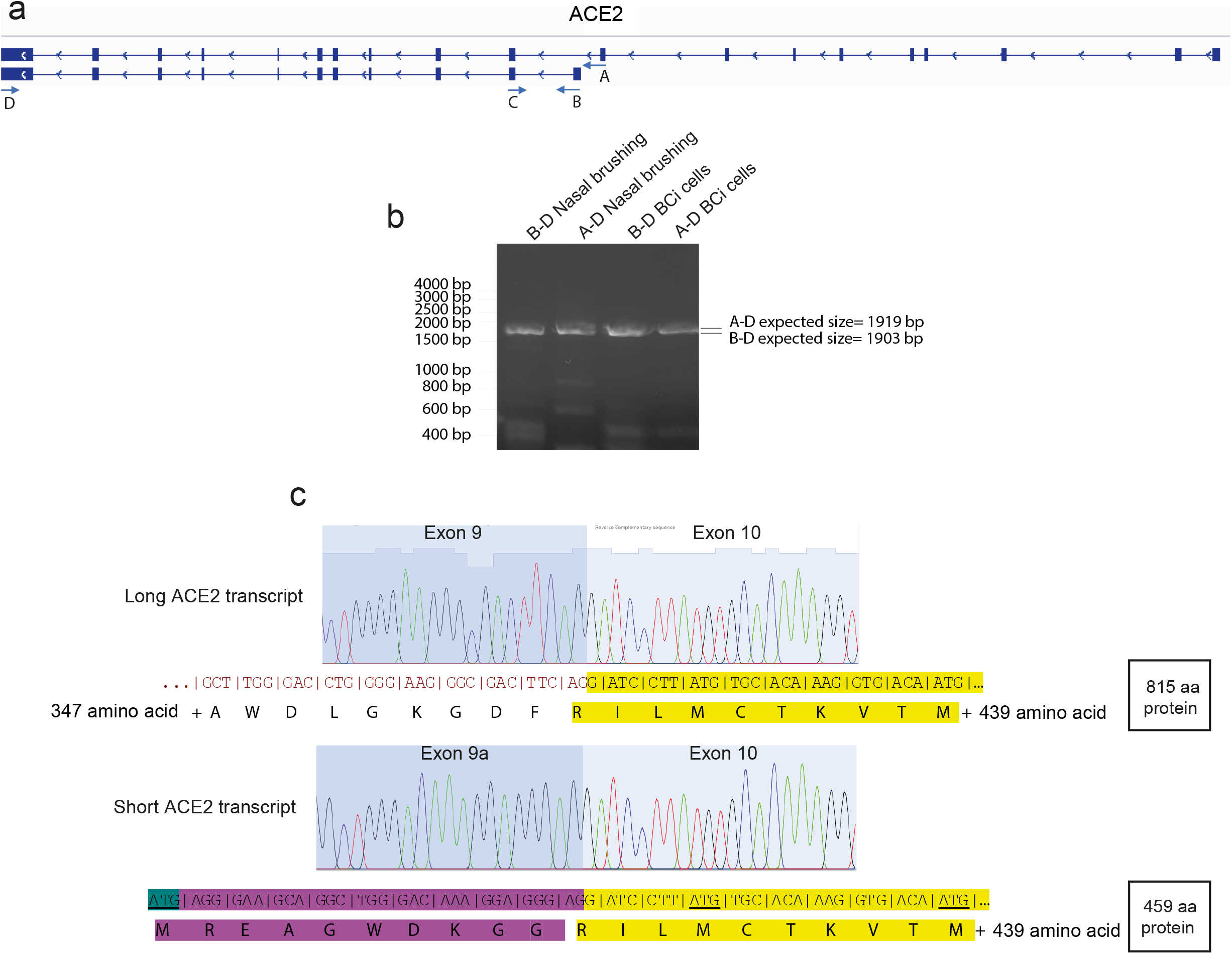
RT-PCR and Sanger sequencing confirm short *ACE2* transcript expression in nasal and bronchial epithelial cells. 2a. Graphic of ACE2 transcripts and RT-PCR primer locations 2b. Agarose gel electrophoresis image of long-range transcript-specific PCR products amplifying full short ACE2 transcript and exons 9-19 of long ACE2 transcript from nasal epithelial brushings and BCi-NS1.1 cells 2c. Sanger sequencing electropherogram traces showing sequence at exon/exon boundaries of long ACE2 transcript exon 9-10 and short ACE2 transcript exon 9a-10. Amino acid translation is shown below

To investigate expression of this novel *ACE2* transcript relative to full-length *ACE2* transcripts (ACE-202 and ACE-201) we extracted the number of reads mapped to exon9a/exon10 junction and reads mapping to exon9/exon10 from the STAR alignment output file and calculated exon 9a inclusion rates relative to inclusion of exon 9 using the following calculation: PSI_new = reads[9a to 10]/(reads[9a to 10] + reads[9 to 10]). This analysis showed that in nasal epithelia the mean expression level of short and long *ACE2* transcripts was 0.745 (reads mapped to exon9a/10 or exon9/10 per million mapped reads) and relative inclusion of exon 9a was 0.763 (st err 0.0829). This is a statistically significant higher level of expression of short *ACE2* than long *ACE2* in nasal epithelial cells (p<0.05, Student’s t-test, n=6). We then designed specific qPCR primers to amplify the short and long transcripts of *ACE2* individually, as well as a pair of common primers to amplify both transcripts and quantify total levels of *ACE2* expression. Expression of long *ACE2* was confirmed in a number of cell lines and primary airway cells, with highest expression being observed in the Caco2 cell line and nasal epithelial cells grown at ALI whose expression was comparable to that observed *ex vivo* (**Figure 3a,b**). With the exception of Caco2 cells, expression of short *ACE2* was very low in the cell lines studied, and this contrasted with the airway cells which exhibited high expression of this novel isoform (**Figure 3a,b**). As we observed high expression of both *ACE2* isoforms in differentiated airway cultures, we assessed its induction during differentiation of nasal epithelial cells grown at ALI *in vitro*. We observed very low *ACE2* expression on day 0, with expression reaching levels comparable with those observed in primary nasal brushings at day 4 of ALI culture when the early start of cilia gene transcription is observed (usually days 4-7), being maintained until day 37, and reducing at day 84 as the cultures became senescent (**Figure 3c and Supplementary Figure 1**). This is consistent with published work showing that *ACE2* expression (and SARS-CoV-2 infection) is dependent on airway epithelial cell differentiation^38^. At all time-points the level of short *ACE2* transcript expression was higher than expression of the long *ACE2* transcripts, although this was not statistically significant. Given reports that bronchial epithelial cells express lower levels of *ACE2* than nasal cells^26^, we also compared expression of the long and short isoforms of *ACE2* in these two cell types. Consistent with previous reports, total levels of *ACE2* were lower in bronchial epithelial cells, which was due to reduced expression of both long and short forms of *ACE2* (**Figure 3d**).

**Figure 3.**
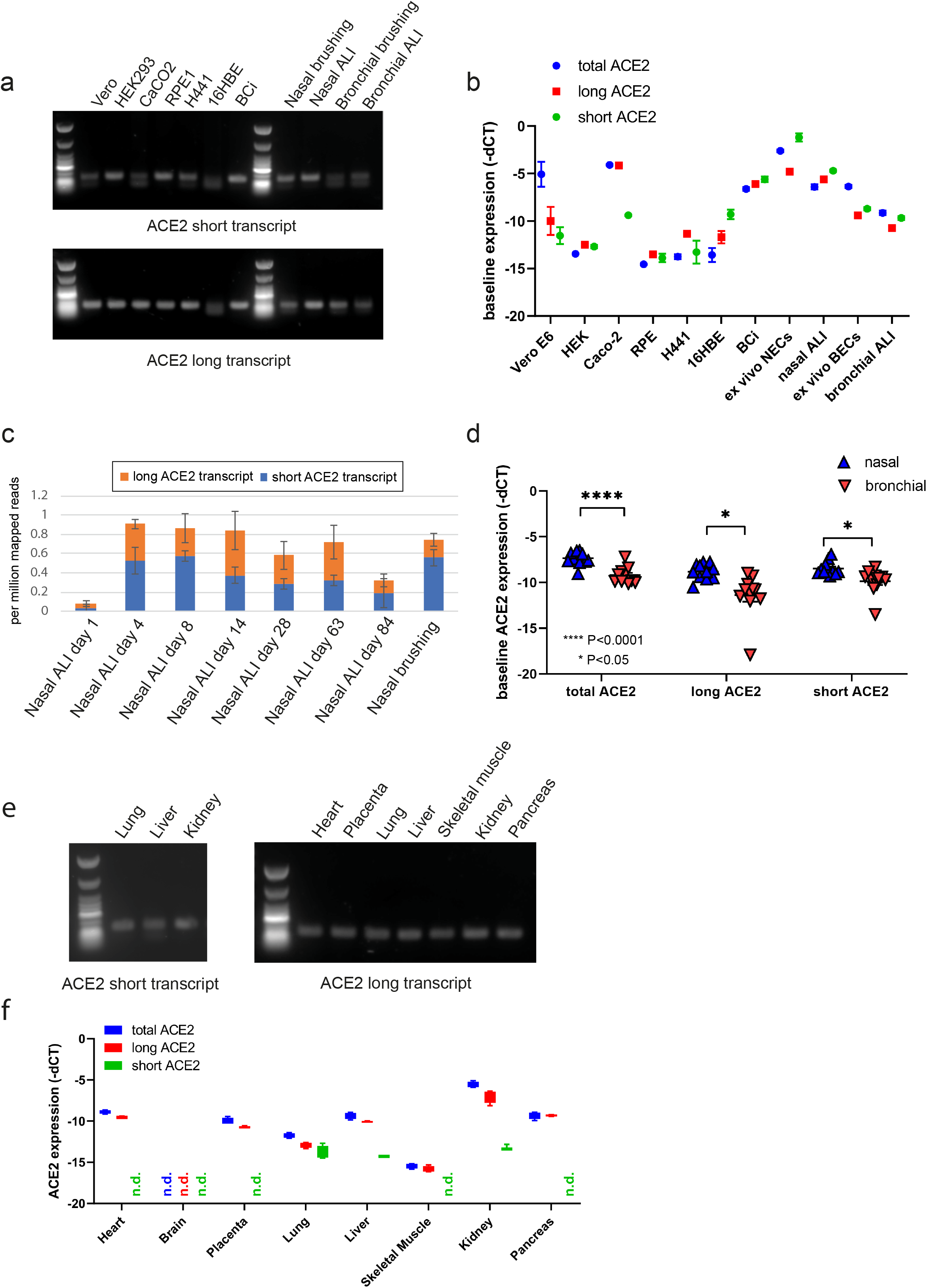
Short *ACE2* is expressed in different cell types and lung, liver and kidney. 3a. Agarose gel electrophoresis image of transcript-specific ACE2 RT-PCRs from different cell types studied. Size standard = NEB Low Molecular Weight ladder. 3b. RT-qPCR analysis of transcript-specific PCRs for long and short ACE2 expression in cell lines and airway cells. Analysis was done at least in duplicate and from different passages or donors in all lines but RPE-1, Caco-2 and HEK 293 cells where analysis was done in duplicate from one passage. 3c. Graphs showing relative expression of short ACE2 transcript and long ACE2 transcript in nasal epithelial cells at different stages of differentiation at air-liquid interface (Day 1, 4, 8, 14, week 4, week 9, week 12) (n=3 for each time point except week 12 where n=2) and primary nasal brushings (n=6). Scale = reads mapped to exon/exon boundary per million mapped reads. Error bars = standard error of the mean. 3d. Graphs showing relative expression of short *ACE2* transcript and long ACE2 transcript in nasal (n=11) and bronchial (n=11) ALI cultures from healthy donors, as determined using transcript specific qPCR. Data were analysed using Mann Whitney U test. 3e. Agarose gel electrophoresis image of transcript-specific PCR products multiple cell types on multiple tissue control panel. This shows expression of the long transcript of *ACE2* in heart, placenta lung, liver skeletal muscle, kidney and pancreas but expression of the short transcript of *ACE2* only in lung, liver and kidney. 3f. Graphs showing relative expression of short *ACE2*, long *ACE2* and total *ACE2* transcript in a Multiple Tissue cDNA panel 1 (636742, Takara), as determined using transcript-specific *ACE2* RT-qPCR. n.d.= not detected. Analysis was done in duplicate runs on the same day, n=2 in different days.

### Short ACE2 transcript is expressed in lung, liver and kidney

We then investigated whether the shorter transcript of *ACE2* is expressed in tissues outside the airway by performing transcript-specific qPCR on cDNA from a multiple tissue control panel. This showed expression of the long transcript of *ACE2* in all tissues tested except whole brain (**Figure 3e,f**), however the short transcript of *ACE2* was only detected robustly in lung, liver and kidney (**Figure 3e,f**) and was below detection within the dynamic range of the qPCR (**Supplementary Figure 2**) in all other tissue types tested. In the lung, short and long ACE2 are expressed at approximately equivalent levels, but in liver and kidney long ACE2 is expressed at a higher level than short ACE2 (**Figure 3e,f**). Together these data show that we identified a novel *ACE2* transcript that is expressed in liver, kidney, and particularly in the lung and airway suggesting a significant role in this compartment.

### Short ACE2 encodes a novel protein lacking most of SARS-CoV-2 binding interface but retains domains required for host protease cleavage

Having confirmed expression of this novel transcript in multiple cell types we sought to investigate whether this transcript is translated into a protein product. Having identified a TATA box and in-frame ATG start codon in the new exon we predicted that this novel transcript would produce a 459 amino acid protein consisting of Arg357 – Phe805 of the full-length long ACE2 protein isoform plus an additional 10 novel amino acids (aa) before Arg357 (M-R-E-A-G-W-D-K-G-G). This is predicted to produce a protein of 52.7kDa which includes the C-terminal 449 amino acids of long ACE2, but lacks the 356 N-terminal residues. Consistent with our expectations, data mining of proteomics datasets in the public domain also identified sequences corresponding to the novel 10 amino acid peptide at the N-terminus of short ACE2 (M-R-E-A-G-W-D-K-G-G; M-R-E-A-G-W-D-K) in colon, breast and ovarian cancer proteomes^39^ therefore supporting protein expression studies.

To investigate whether an ACE2 protein isoform of the expected size is expressed in airway epithelial cells and other cell types, we performed western blotting cells using multiple antibodies to ACE2 recognising epitopes on different regions of the protein (**Figure 4a**). We anticipated that short ACE2 would be detected by antibodies raised to the C-terminal domain (CTD) of ACE2 but not the N-terminal domain (NTD) of ACE2. Using a CTD ACE2 antibody (Abcam 15348), with Vero cells as a control, we observed two bands at 100 and 120kDa, consistent with the presence of glycosylated and non-glycosylated forms of full length ACE2 protein^14^. In nasal epithelial cells, we also observed these two isoforms, and an additional band at ~50kDa, the expected molecular weight of short ACE2. (**Figure 4b**). Furthermore, this 50kDa band was at least as intense as full length ACE2, consistent with our qPCR analyses of nasal epithelium. To orthogonally validate antibody specificity, we used an antibody raised against the NTD of ACE2 (Abcam 108252) in nasal epithelial cells (**Figure 4b**). Our data show that the 50kDa band is only recognized by the antibody raised against anti-ACE2 CTD domain antibody and not by anti-ACE2 NTD domain antibody as expected. Using a validated ACE2 antibody which recognises epitopes from aa 18-740 (R&D AF933) we also observed the 50kDa band in bronchial epithelial cells grown at ALI (**Figure 4b**).

**Figure 4.**
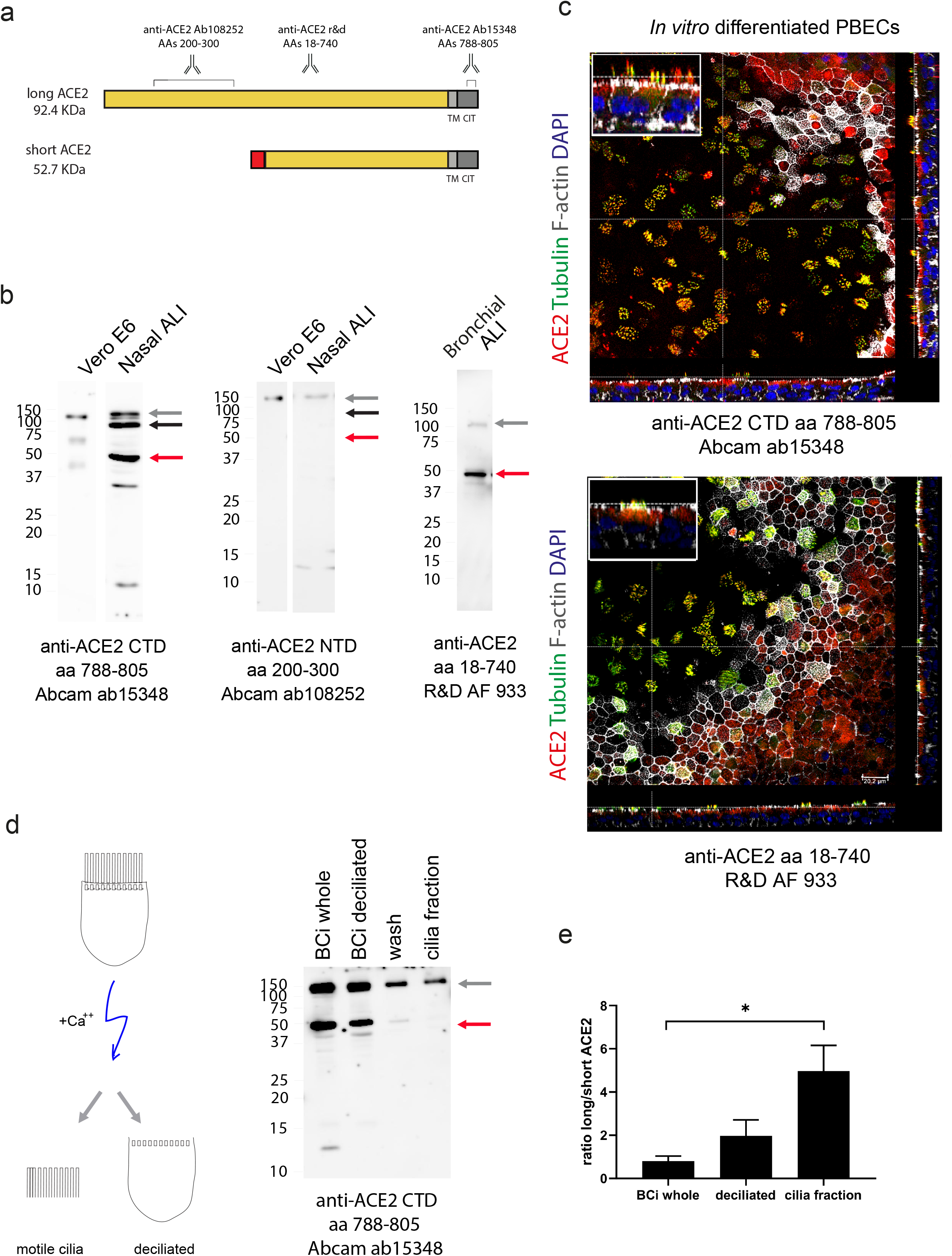
Short ACE2 protein is expressed and is not enriched on motile cilia relative to long ACE2 protein. 4a. Schematic illustration of predicted long and short protein isoforms of ACE2 and position of antigen sequences used to generate antibodies used. 4b. Left panel: Representative western blot of Vero cells and nasal cells (n=3) grown at ALI blotted with ACE2 antibody raised to C-terminal domain (amino acids 788-805). Grey arrow points to glycosylated long ACE2 detected in both Vero E6 and nasal ALIs, black arrow points to unglycosylated long ACE2 in nasal ALI, red arrow points to unglycosylated short ACE2 in nasal ALI. Middle panel: Representative western blot of Vero cells and nasal cells grown at ALI blotted (n=3) with ACE2 antibody raised to N-terminal domain (amino acids 200-300). Grey arrow points to glycosylated long ACE2 detected in both cell types. Right panel: Western blot of bronchial cells grown at ALI blotted with ACE2 antibody raised to epitopes across the protein (amino acids 18-740). 4c. Representative IF confocal images of ALI differentiated primary bronchial epithelial cells stained with anti-alpha tubulin (green), Acti-Stain 555 Phalloidin (grey), DAPI (blue) and ACE2 (red) detected with C-terminal domain antibody (top panel) or antibody detecting epitopes across the protein (bottom panel). N=2 4d. Schematic illustration of deciliation protocol using calcium shock (left) and western blot of whole and deciliated BCi-NS1.1 cells, deciliation wash, and cilia pellet (right). Red arrow points to short ACE2 which not enriched on cilia relative to long ACE2 enrichment. The graph shows semi-quantitative analysis of the Western blots by densitometric analysis (n=4).

To examine the localisation of the ACE2 isoforms in airway epithelial cells, we performed immunofluorescent staining of differentiated ALI cultures of primary bronchial epithelial cells with anti-ACE2 antibodies visualised by confocal microscopy (**Figure 4c, Supplementary Figure 3**). The antibodies which recognise common epitopes in short and long ACE2 localised mainly to the apical portions of the cells and to motile cilia. Interestingly, ACE2 staining extended further up the cilium than the microtubular axoneme, giving the impression of staining the ciliary tip, whereas, in fact, it stained the full length (**Figure 4c, Supplementary Figure 3**). Staining with the third antibody that detects only long ACE2 was too weak to interpret. However, staining with the third antibody that detected only long ACE2 was too weak to interpret.

Thus, to further investigate whether short ACE2 localises to cilia as has been reported for full-length ACE2^40^ we extracted total protein from BCi-NS1.1 cells hTERT immortalized bronchial cells which differentiate robustly into airway multiciliated cells^37^, deciliated BCi-NS1.1 cells and cilia purified from BCi-NS1.1 cells and analysed by western blot with antibodies specific to the C-terminus of full-length ACE2 (Ab15348, aa788-805). Consistent with our previous experiments, this analysis showed a distinct band around 50kDa detected by the C-terminal ACE2 antibody (**Figure 4d**). Notably, the band corresponding to short ACE2 was not enriched in the cilia fraction albeit still present in detectable amounts (**Figure 4d**) suggesting that it is predominantly localised to the apical membranes of the cells. Densitometry analysis of western blot images confirmed enrichment of long ACE2 in cilia fraction relative to short ACE2 (**Figure 4d**). Furthermore, long and short ACE2 were both found in the deciliated cells in a ratio of approximately 1:1 - 2:1, suggesting the possibility of heterodimer formation between the two isoforms.

### Novel short ACE2 isoform forms a stable dimer lacking most of SARS-CoV-2 binding interface

To better understand the possible functions of the short isoform of ACE2 we modelled the structure of this short isoform, based on the full-length structure of long ACE2 resolved by cryo-EM (PDB 6M18)^17^. This analysis highlighted the extent of loss of the SARS-CoV-2 binding region in short ACE2, with many residues previously shown to be important for viral binding not present in short ACE2 sequence (**Figure 5a-c**)^17^. In particular, short ACE2 lacks two entire regions involved in interaction with SARS-CoV-2 spike glycoprotein (aa 30-41 and aa 82-84), including a high-affinity binding site (aa 30-41), but retains part of a third region involved in this interaction^17^. This latter sequence is replaced by the N-terminal specific sequence of short ACE2, which is predicted to form a disordered/helical secondary structure by PEP-fold, compared to the beta sheet present in long ACE2 further modifying the third binding interface to Spike. Short ACE2 however retains the sequences required for cleavage by ADAM17, TMPRSS11D and TMPRSS2, suggesting that can be a substrate of these proteases. Interestingly, the ACE2 residue critical for substrate selectivity to Angiotensin II, Arg514, is missing from short ACE2 suggesting that it may lack catalytic activity toward Angiotensin II. Altogether, this analysis suggests that short ACE is not competent for high affinity binding to SARS-CoV-2 spike protein, but it can be a substrate of host proteases acting on long ACE2 during viral entry.

**Figure 5.**
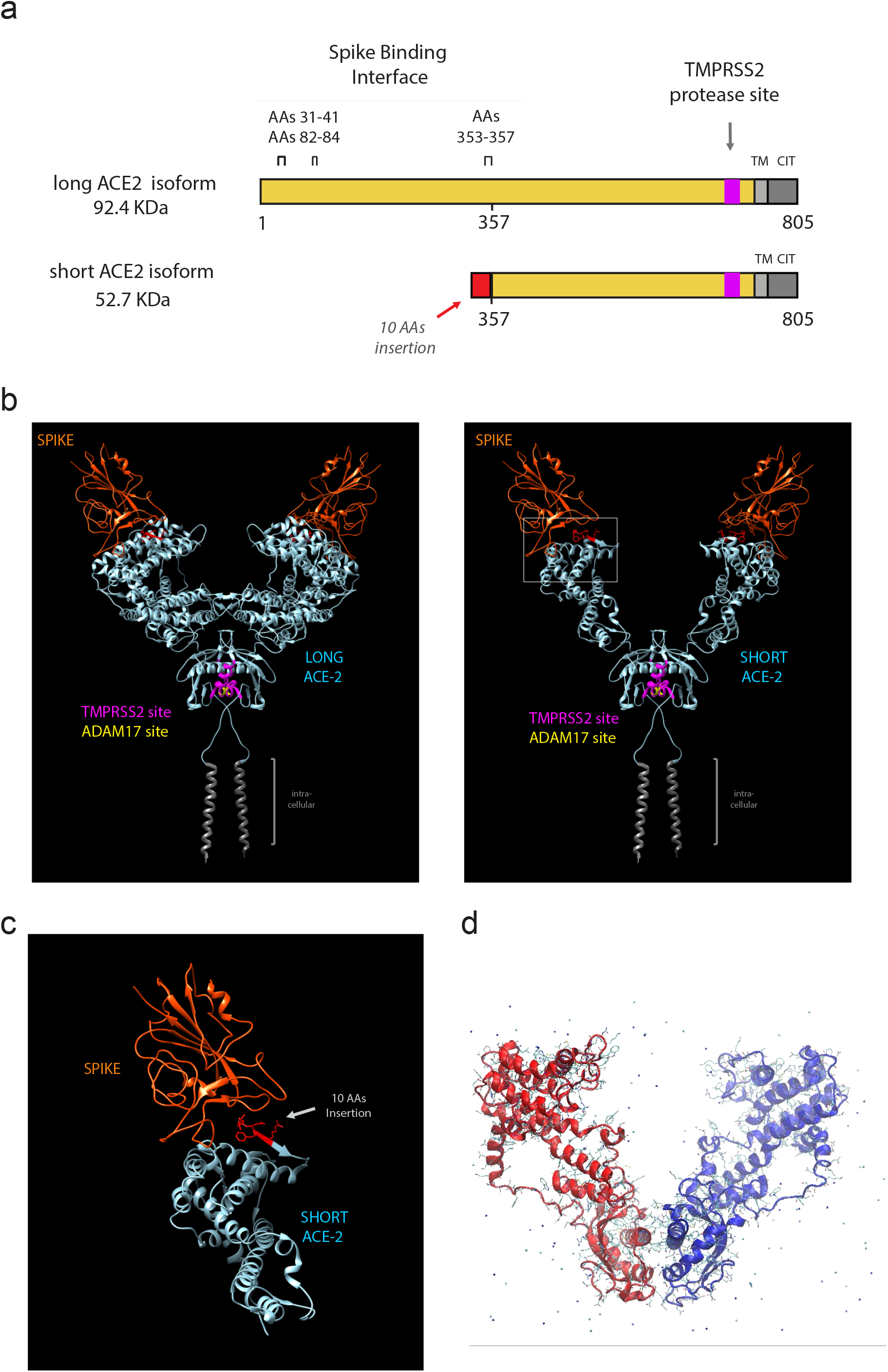
Short ACE2 lacks most of the Spike binding regions but contains TMPRSS2 protease site. 5a. Primary sequence of long and short ACE2 with position of relevant spike binding regions and protease cleavage domains. 5b. Long ACE2 or short ACE2 (from pdb 6M18, teal) in complex with SARS-Cov-2 spike protein (orange). Residues essential for cleavage by ADAM17 are shown in yellow and residues essential for cleavage by TMPRSS11D and TMPRSS2 are shown in fuchsia. 5c. High magnification of the putative interaction region of short ACE2 with virus spike protein. 5d. Snapshot of MD simulation of short ACE2 obtained with YASARA (**Supplementary Video 1**)

While the majority of the spike binding domain of ACE2 is missing from short ACE2, the neck domain (also called the ferredoxin-like fold domain (residues 616 to 726) which is the most important dimerization interface^17^ and transmembrane region are present in short ACE2. This suggests that ACE2 may exist not only as the conventional full length ACE2 homodimer, but also as a short ACE2 homodimer and a heterodimer of full length ACE2 and short ACE2 (**Supplementary Figure 4**).

Since a substantial portion of ACE2 is missing in short ACE2, we then undertook molecular dynamic simulation of short ACE2 dimer to assess its dimerization and stability. Assuming that the short form folds in the homologous parts in the same way as the full length ACE2, molecular dynamic simulation of short ACE2 dimer and analysis of secondary structure elements changes show that this structure is likely a stable dimer, as suggested by the retention of the sequence corresponding to the Neck dimerization domain, the main dimerization interface between the monomers^17^ (**Figure 5d, Supplemental Figure 4 and Supplementary Video 1**).

### Novel short ACE2 isoform is upregulated by interferon and rhinovirus (RV) infection

To begin investigating the functional relevance of short *ACE2* transcript expression, we first assessed whether it was an IFN stimulated gene. As expected, treatment of bronchial epithelial cells with IFN-beta resulted in upregulation of the IFN-responsive genes, MxA and IP10. We also observed upregulation of total *ACE2*, which to our surprise was largely due to an effect on short *ACE2* rather than long *ACE2* (**Figure 6a**). This led us to evaluate the response to viral infection, as it has been reported that *ACE2* expression is upregulated in this condition^26^. We exposed nasal epithelial cells grown at ALI to rhinovirus (RV) and harvested cells 24hr after infection. qPCR analysis showed a significant upregulation of both long and short *ACE2* RNA expression relative to UV-RV treated control (*: p=<0.001, non-parametric Wilcoxon Signed Rank test, control vs. RV16 n=11) (**Figure 6b**). While this is consistent with previously published work showing that *ACE2* is upregulated in response to influenza exposure^26^, it is notable that we found that it was the short *ACE2* transcript which was upregulated more robustly than long *ACE2* transcript (around 9-fold increase in expression of short *ACE2* compared to around 2.5-fold increase in long *ACE2* expression) (**Figure 6b**). Parallel experiments using bronchial epithelial cells infected with RV, confirmed induction of short *ACE2* but no significant effect on long *ACE2* (**Figure 6b**).

**Figure 6.**
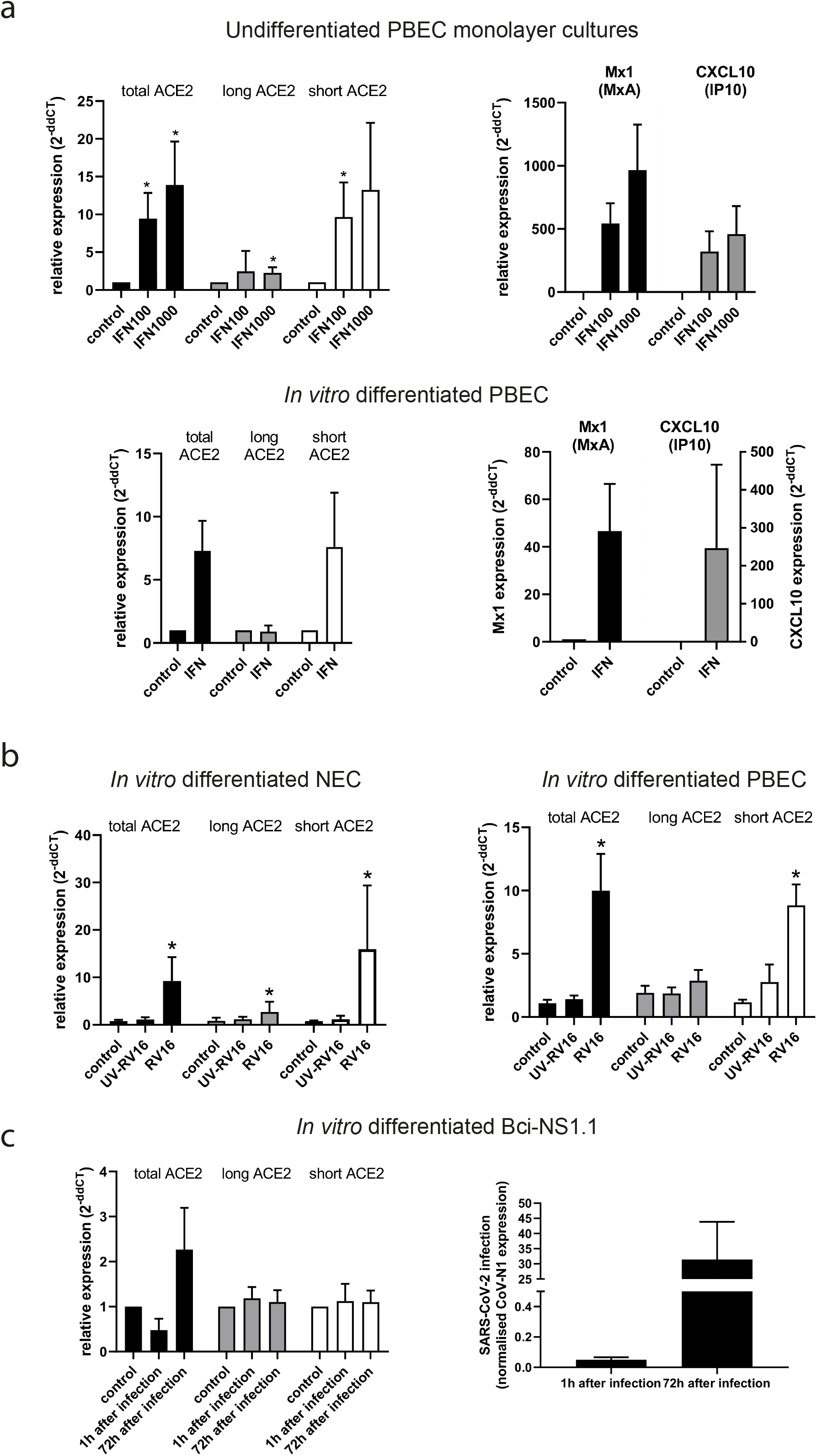
Short *ACE2* is upregulated in response to IFN-beta and rhinovirus (RV16) infection but not SARS-CoV-2 infection. 6a. Undifferentiated primary bronchial epithelial cell (PBEC) monolayer cultures (N=3) (top) or *in vitro* differentiated (ALI) PBEC cultures (N=3) (bottom) were treated with IFN-beta (100 or 1000 IU/ml) for 24h and ACE2 transcripts (left panel) and induction of IFN-response genes (MX1 and CXCL10) (right panel) were measured by RT-qPCR. Data were analysed using Students t-test. 6b. *in vitro* differentiated (ALI) nasal epithelia cells (NEC) (n=11) (left panel) or *in vitro* differentiated (ALI) bronchial epithelia cells (BEC) (n=11) (right panel) were infected with rhinovirus (RV16) (MOI of 1) or mock-infected using a UV-irradiated control (UV-RV16). Nasal cells were collected from 3 female, 8 male patients with a mean age of 45.31+/-3.23 (SEM). After 24h, induction of *ACE2* isoform expression was assessed by RT-qPCR with transcript-specific primers. Data were analysed using non-parametric Wilcoxon test. 6c. BCi-NS1.1 cells were grown at ALI and then infected on the apical side for 1 hour with 100,000 pfu of SARS-Cov-2 strain nCoV/Victoria/1/2020 obtained from Public Health England (PHE), UK. Cells were harvested in QIAzol at 1h post infection and at 72h, RNA extracted and quantitative RT-qPCR performed to detect SARS-CoV-2 using 2019-nCoV_N1 primers and the housekeeping genes HPRT, 18S and RNAse P using the dCt method. 1 hour and 72 hours after infection induction of *ACE2* transcript expression was assessed by RT-qPCR with transcript-specific primers (left). SARS-CoV-1 infection was confirmed by CoV-N1 RT-qPCR 1 and 72 hours after infection (right). Data were analysed using non-parametric Wilcoxon test. N=4.

As long ACE2 has been described as a point of entry for SARS-CoV-2, and that SARS-CoV-2 infection stimulates an increase in *ACE2* expression, we sought to investigate the effect of SARS-CoV-2 infection on the expression of the short transcript of *ACE2*. As infection model we used BCi-NS1.1. BCi-NS1.1 were infected with SARS-CoV-2 and harvested cells at 1 and 72 hours after infection. qPCR amplification of the CoVN1 nucleocapsid gene expression confirmed significant infection by 72 hours (**Figure 6c**), however we failed to see significant induction of either the long or short isoforms of *ACE2*, or total *ACE2* (**Figure 6c**). As *ACE2* isoforms are both inducible by IFN, it is likely that lack of *ACE2* induction at this time point is due to the known ability of SARS coronaviruses including SARS-CoV-2 to inhibit both IFN expression and downstream signalling from the Type I IFN receptor^41,42^.

### Short and long ACE2 are regulated differently in severe asthma

It has been reported that patients with asthma have reduced susceptibility to SARS-CoV-2, and that asthma symptoms are not exacerbated by SARS-CoV-2 infection^43–45^. However, it has been suggested that a subset of asthma patients (Type-2 low asthmatics) who are more at risk of SARS-CoV-2 show higher *ACE2* expression in bronchial epithelium, associated with upregulation of viral response genes^46^.

To explore whether expression of short *ACE2* might have a role in SARS-CoV-2 infection in asthma, we investigated expression of the long and short transcripts of *ACE2*. Transcript-specific qPCR on cDNA from bronchial brushings from healthy controls and patients with severe asthma showed statistically significantly lower expression of total *ACE2* and long *ACE2* in patients with severe asthma, but no significant difference in short *ACE2* expression between groups (**Figure 7a**). This difference in the profile of *ACE2* expression may be a feature of the disease itself, an effect of treatment, or a combination of both. For example, in asthma there is goblet cell metaplasia resulting in a reduction in the number of ciliated cells^47^ which are the site of *ACE2* expression while elevated levels of the type 2 cytokine, IL-13, are reported to suppress *ACE2* expression^48^. Since all of the severe asthmatic subjects received inhaled or oral corticosteroids as part of the regular care, it is noteworthy that oral or intravenous dexamethasone treatment of mechanically ventilated Covid-19 patients resulted in improved 28-day mortality^49^. While corticosteroids are well known as anti-inflammatory agents, whether they also directly modulate *ACE2* expression is unknown.

**Figure 7.**
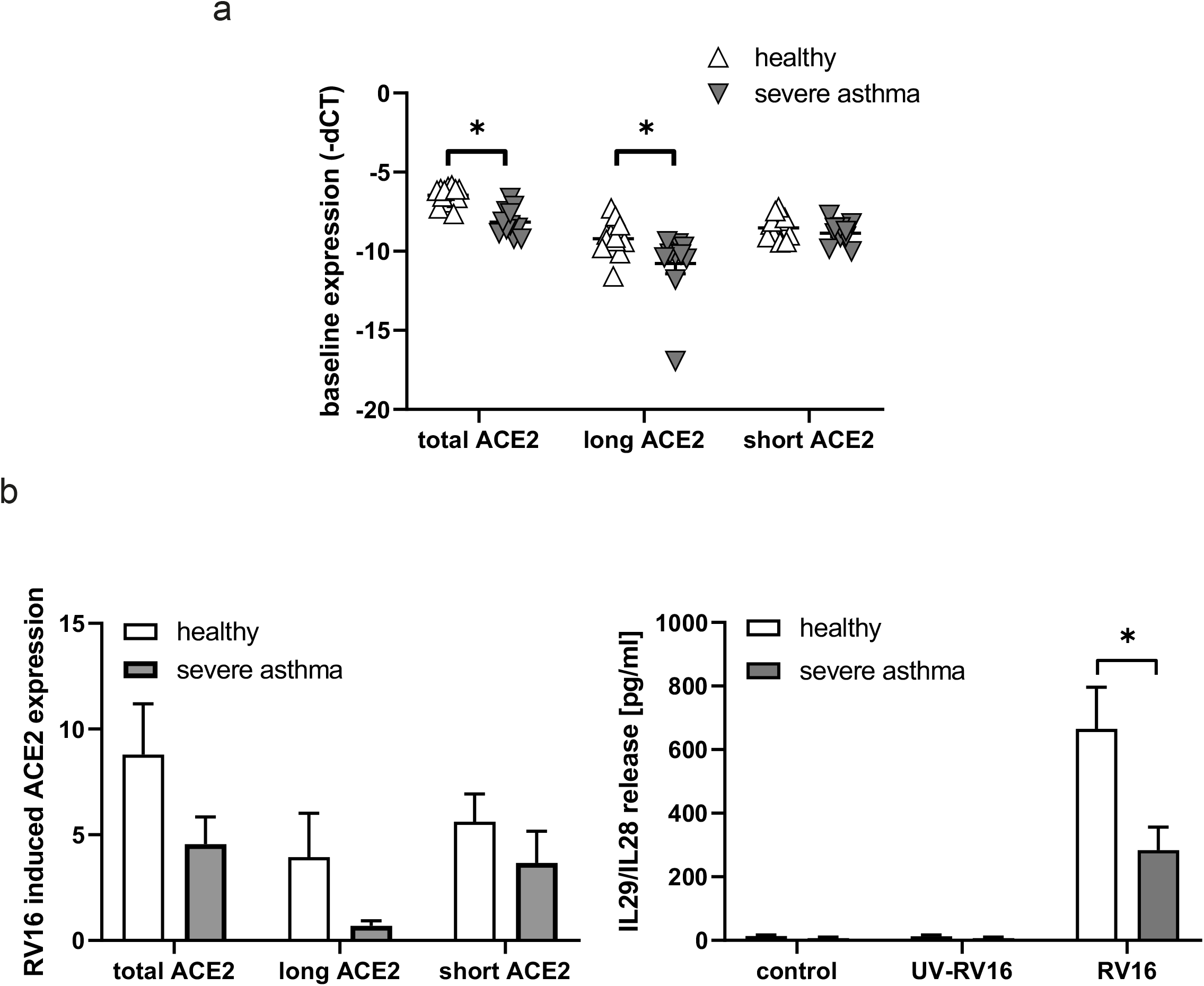
Long *ACE2* is expressed at lower levels in bronchial brushings from severe asthmatic donors and RV-induced expression of long *ACE2* is reduced in bronchial ALI cultures from severe asthmatic donors. 7a. Bronchial epithelial brushes from healthy controls (n=13) or severe asthmatic (n=11) donors were harvested and RNA extracted for analysis of *ACE2* transcript expression by transcript-specific RT-qPCR. Data were analysed using nonparametric Mann-Whitney test. 7b. Bronchial epithelial cells from healthy (n=11) or severe asthmatic (n=7) donors were grown at ALI and infected with rhinovirus (RV16) (MOI of 1) or mock-infected using a UV-irradiated control. After 24h, induction of *ACE2* transcripts was assessed by RT-qPCR with transcript-specific primers in duplicate. The fold-induction of each transcript by RV16 was quantified using the ddCt method using UV-RV exposed cells as control. Activation of anti-viral response in RV16-infected ALI cultures was demonstrated by detection of IL29/IL28 in basolateral supernatants by ELISA (healthy n=14, severe n=8). Data were analysed using Student’s t-test.

Asthma patients have also been shown to mount a reduced antiviral IFN response following infection with respiratory viruses resulting in disease exacerbation with a risk of hospitalisation or even death^50–55^. As we observed that short *ACE2* was not induced following SARS-CoV-2 infection where IFN expression and signalling are compromised, we examined its induction in differentiated bronchial epithelial cultures from severe asthmatic donors in response to RV infection. When compared to cultures from healthy control donors, bronchial epithelial cells from patients with severe asthma showed less upregulation of total *ACE2* and long *ACE2*, whilst short *ACE2* was induced to almost the same levels as in control cultures. (**Figure 5c**). As we observed previously^54^, IFN lambda was also reduced in cultures from severe asthmatic donors. Together, these data are consistent with a hypothesis that suppression of long *ACE2*, alongside maintenance of short *ACE2* expression is protective against SARS-CoV-2 infection in severe asthmatics.

## Discussion

Here we present identification and characterisation of a short 11 exon transcript of human *ACE2*, consisting of a novel previously unannotated first exon, which we name exon 9a, and exons 10-19 of the long *ACE2* transcript. We show that, whilst the long transcript of *ACE2* is expressed in multiple tissues, this short ACE2 transcript is expressed exclusively in airways, liver and kidney. We show highest expression in primary respiratory epithelia, most notably in the nasal epithelium where the level of short transcript expression is higher than long *ACE2* transcript expression. Expression of both transcripts of *ACE2* is dependent on differentiation of epithelial cells, with levels of expression comparable with primary nasal epithelia from days 4 – 63 of differentiation at ALI culture. We show that transcription of this short transcript is regulated independently of the long transcript of *ACE2*, with putative promoter elements identified upstream of the transcriptional start site of the short *ACE2* transcript. We confirm that this short transcript is translated into a protein product of around 52kDa, a 459 amino acid isoform of ACE2 which lacks a signal peptide. Modelling suggests that this isoform contains a transmembrane domain, collectrin homology domain and portions of the ACE homology domain but lacks the 356 most N-terminal amino acids of full-length ACE2, with 10 novel amino acids in their place. Most of the key residues required for SARS-CoV-2 spike binding are absent from this isoform and we hypothesise that short ACE2 is not a viral entry point for SARS-CoV-2. Molecular dynamic simulation suggests that this short isoform can form thermodynamically stable dimers, but the function of this short form of ACE2 remains unclear. Given that the short isoform of ACE2 retains its transmembrane domain and collectrin domain it seems possible that the short isoform may be functionally involved in regulation of amino acid transport in the airway. The short form also retains the catalytic residues conferring carboxypeptidase function, suggesting that the short isoform may retain some catalytic activity, albeit likely with different substrate specificity. Structural studies of full-length ACE2 catalytic domain suggest that lack of the N-terminal residues of ACE2 (especially R273) may affect substrate specificity, suggesting that angiotensin-II may not be the primary substrate of the short ACE2 isoform. We further show that the short transcript is an interferon-regulated gene and is more strongly induced by IFN-beta and viral infection than long *ACE2*.

Although the function of the short *ACE2* isoform remains unclear, our data clearly show that this isoform is expressed in the airways, particularly in ciliated cells of the nasal and bronchial epithelium. Of note, ACE2 is an homologue of ACE which also utilizes different promoters to produce two distinct isoforms, the full length molecule and a lower molecular mass variant which is found only in testis (tACE)^56^ where its expression regulates sperm capacitation^57^. Like the motile cilia of the airways, sperm flagella have a characteristic 9+2 axoneme structure suggesting some important function of the shorter ACE2 and ACE isoforms, respectively, in relation to these structures. However, we did not find evidence that short ACE2 was enriched in cilia which contrasts with the long form of ACE2. Instead, we found that short ACE2 was retained within the cell body along with a fraction of long ACE2. Therefore, it is interesting that short ACE2 is preferentially upregulated in response to viral infection, independently of long transcript *ACE2* expression. We hypothesise that this short isoform of ACE2 plays an important physiological role in the airway and, in addition that it may influence host susceptibility to SARS-CoV-2 infection either by heterodimerising with long ACE2 to prevent its trafficking to the exposed tips of the cilia and/or influencing the spike binding interaction; alternatively it may compete for other membrane proteases required for viral entry.

This finding may have significant consequences for design of therapeutic approaches to tackling COVID-19. At present, therapeutic approaches in development include neutralising antibodies designed to bind spike receptor binding domain (RBD) with higher affinity than recombinant human ACE2. These strategies use the extracellular domain of recombinant human ACE2 to design peptides to bind to spike RBD. This assumes full length ACE2 forms homodimers, and the discovery of short ACE2 and its potential ability to form heterodimers may have important impacts on drug design^58^. Other strategies aim to use purified soluble recombinant ACE2, or ACE2+-small extracellular vesicles^59^ to ‘intercept’ the virus before entry into host cells, avoiding infection. Our discovery also has implications for the numerous studies reporting on ACE2 expression levels and differences in levels of expression along airways, across age groups and disease groups^29,46,60,61^ including COVID-19 disease severity^61,62^. This finding should be considered in in choice of reagents for future studies of *ACE2* expression in tissues relevant to SARS-CoV-2 viral infection, bearing in mind the genomic region targeted by primer sets and the epitopes recognised by antibodies. For example, this may help to resolve the current debate over the relevance of conjunctiva as a site of SARS-CoV-2 viral entry, with IHC and protein studies suggesting expression of ACE2 but some RNA studies disputing this^63–66^.

Finally, this data emphasises the remaining gaps in our understanding of full complexity of the human transcriptome, particularly cell-specific transcriptomes, and the power of transcript-level analysis of deep bulk RNA sequence data in resolving some of this complexity. A recent landmark study of unannotated transcripts across 42 different human tissues highlighted the major gaps in transcript annotation of even very intensively studied disease-relevant genes, and the impact this has on our understanding of disease^67^. Whilst advances in RNA sequencing technologies have seen a recent increase in RNA expression studies, many studies focus on gene-level expression analyses, and our understanding of transcript-level expression remains poor. In particular, the limited depth of single cell RNA sequencing does not permit transcript-level analysis, and as a result this is neglected in much cell sequencing data. Deep, bulk RNA sequence analysis of isolated cell types followed by transcript level analysis, including transcript expression quantification and differential splicing analysis has the power to revolutionise understanding of disease.

## Materials and Methods

### Collection of airway samples

Nasal epithelial cells were isolated by brushing the inferior turbinate with a sterile 3.0 mm cytology brush (Conmed). Cells were processed into RNAlater for subsequent RNAseq analysis or were stored in liquid nitrogen prior to cell culture. Bronchial epithelial cells were harvested by bronchoscopic brushings for primary bronchial epithelial cell culture. Airway samples for the study were collected following approval by South-Central Hampshire A, Research Ethics Committee, UK (reference numbers: 07/Q1702/109, 13/SC/0182 and 14/WM/1226) and all participants gave their informed consent.

### Cell culture

Nasal cells were expanded and differentiated at air-liquid interface (ALI) culture as previously described^68^. All cultureware were pre-coated in 1:10 diluted PureCol collagen (5005-B CellSystems) throughout each step. Cells were cultured using Pneumacult Ex-Plus (Stemcell Technologies) and at ALI on 6.5mm 0.4μm polyester membrane transwell permeable supports (Corning Life Sciences) for up to 84 days using Pneumacult ALI media (Stemcell Technologies), with apical surface washed (HBSS) and medium changes 3 times weekly. Cell were cultured in 100% relative humidity, 5% CO_2_ at 37°C. First ciliation was observed by microscopy from day 7 at ALI and maintained until harvested. Primary nasal epithelial cells were fully differentiated and ciliated by 28 days.

Primary bronchial epithelial cells were expanded in Airway Epithelial Cell Growth medium (Promocell) up to passage 1 as previously described^69^. At passage 2 cells were either cultured submerged as monolayers or differentiation was induced by plating cells on 6.5mm 0.4μm polyester membrane transwell permeable supports (Corning Life Sciences) and differentiated at ALI for 21 days. Transepithelial electrical resistance was monitored weekly using an EVOM Voltohmmeter (World Precision Instruments) and cells with a TER ≥330WΩm2 on day 21 were used for experiments.

hTERT transformed bronchial epithelial cell line BCi-NS1.1, provided by Walters et al^37,70^ were expanded in PneumaCult-Ex Plus Basal Medium (Stem Cell Technologies) supplemented with Pneumacult Ex Plus supplements (Stem Cell Technologies), hydrocortisone, nystatin and penicillin/streptomycin. BCi-NS1.1 cells were grown at air-liquid interface in PneumaCult-ALI Basal Medium (Stem Cell Technologies) supplemented with Pneumacult ALI supplement (Stem Cell Technologies), hydrocortisone, PneumaCult ALI maintenance supplement, heparin, nystatin and penicillin/streptomycin. BCi-NS1.1 cell ciliation was observed by microscopy and cells were fully differentiated and ciliated by 42 days at ALI.

Vero E6 (ECACC Vero C1008) cells were cultured in DMEM medium supplemented with 10% FBS and penicillin/streptomycin (Gibco) at 37°C with 5% CO_2_. When 70-80% confluent (every 5-7 days) cells were passaged by washing with HBSS before detaching with 0.2% trypsin EDTA.

NCI-H441 [H441] (ATCC HTB-174) cells were cultured in RPMI 1640 medium (Gibco) supplemented with 10% FCS, sodium pyruvate, L-glutamine and penicillin/streptomycin at 37°C with 5% CO_2_ and passaged every 3-4 days. Cells were washed with modified HBSS with calcium and magnesium (HyClone) before detaching with 0.2% trypsin EDTA.

hTERT RPE-1 (ATCC CRL-4000) were cultured in DMEM/F12 medium (Gibco) supplemented with 10% FCS at 37°C with 5% CO_2_ and passaged every 4-6 days. 293 [HEK-293] (ATCC^®^ CRL-1573™) were cultured in DMEM high glucose supplemented with 10% FCS at 5% CO_2_ and passaged every 4-6 days at a ratio of 1:8.

### RNA extraction and quality control for RNAseq

RNA was extracted from samples using RNeasy Plus Mini kit (Qiagen). RNA quality and concentration were measured using an RNA Nano chip on the Bioanalyzer 2100 (Agilent). Samples with total RNA concentration ≥20ng/μl, RIN ≥9.6 and OD 260/280 were taken forward for cDNA library preparation and sequencing.

### cDNA library preparation, sequencing and data quality control

cDNA libraries were prepared using Ribo-Zero Magnetic Kit for rRNA depletion and NEBNext Ultra Directional RNA Library Prep Kit library prep kit by Novogene Inc. Library quality was assessed using a broad range DNA chip on the Agilent Bioanalyser 2100. Library concentration was assessed using Qubit and q-PCR. Libraries were pooled, and paired-end 150bp sequencing to a depth of 20M-100M reads per sample was performed on an Illumina HiSeq2500 by Novogene Inc.

Raw FASTQ reads were subjected to adapter trimming and quality filtering (reads containing N > 10%, reads where >50% of read has Qscore<= 5) by Novogene Inc.

Quality of sequence was assessed using FastQC v0.11.5 (https://www.bioinformatics.babraham.ac.uk/projects/fastqc/). No further data filtering or trimming was applied._Raw FASTQ reads after adapter trimming and quality filtering (reads containing N > 10%, reads where >50% of read has Qscore<= 5) were deposited on the Sequence Read Archive, SRA accession to be provided upon publication.

### Alignment to reference genome and quality control

Paired FASTQ files were aligned to GRCh38 human genome reference using GENCODE v33 gene annotations and STAR v2.6.0a splice aware aligner^31^, using ENCODE recommend options (3.2.2 in the STAR manual (https://github.com/alexdobin/STAR/blob/master/doc/STARmanual.pdf). The two-pass alignment method was used.

Alignment files were assessed for saturation of known splice junctions using RSeqQC v3.0.1^71^.

### Transcriptome assembly

Unique transcripts were assembled from merged alignment files, and a merged transcriptome reference formed from the unique transcripts and GENCODE v33 reference transcriptome using SCALLOP tool v0.10.5^72^.

### Alignment to reference transcriptome and transcript level abundance estimates

SALMON tool v1.3.0^73^ was used to perform transcript abundance estimates from raw FASTQ files using selective alignment with a decoy-aware transcriptome assembled using Scallop tool._Integrative genome viewer (IGV) v.2.3.93^32^ was used to visualise alignment files.

### Differential splicing analysis

A Mendelian RNA-seq method for identifying and filtering splice junctions developed by Cummings *et al*. ^33^ was used to detect aberrant and novel splice events. No changes were made to this code. The individual sample splice junction discovery output files were combined into an overall splice junction discovery file used for splice junction normalisation.

### RNA extraction and cDNA production

<u>From nasal and bronchial brushings</u>, cDNA was synthesised from excess RNA purified for RNAseq using High Capacity cDNA Reverse Transcription kit (Thermo Fisher Scientific) following manufacturer’s instructions.

<u>From cell lines</u>, RNA was isolated from cell lysates using standard phenol-chloroform extraction, and reverse transcribed to cDNA using a Precision Reverse Transcription kit (PrimerDesign, Southampton, UK) according to the manufacturer’s instructions. From BCi-NS1.1, RNA was isolated from cell lysates using standard QIAzol extraction, and reverse transcribed to cDNA using High Capacity cDNA Reverse Transcription kit (Thermo Fisher Scientific) following manufacturer’s instructions.

### Long-range RT-PCR

Phusion High-Fidelity PCR Master Mix with HF Buffer (NEB) was used to amplify the novel and annotated transcript using custom primers (IDT) from cDNA produced from nasal brushings and BCi-NS1.1 cells. Manufacturer’s instructions and recommended thermocycling conditions were followed, with annealing temperature (64°C) calculated using NEB Tm calculator. RT-PCR Primer sequences: For amplifying short ACE transcript (exon 9a - exon 19): forward 5’-3’ ATTGAGGAGAGCTCTGAGGC, reverse 5’-3’ TCTCTCCTTGGCCATGTTGT. For amplifying long ACE2 transcript (exon 1 - exon 19): forward 5’-3’ TGCTAACGGACCCAGGAAAT, reverse 5’-3’ TCTCTCCTTGGCCATGTTGT. Samples were size separated against Hyperladder 1kb (BioLine).

Gel extracted PCR products were sequenced using forward and reverse PCR primers at 3.2**μ**M by Source Biosciences. Electropherograms were visualised using 4Peaks.

### RT-qPCR

cDNA from cell cultures and from human multiple tissues control cDNA panel I (TakaraBio) was amplified by qPCR (cycling conditions 95 °C 10 min, then 50 cycles of 95 °C 15 s, 60 °C 1 min) using ACE2 primer pairs (see below). Data were normalised to the geometric mean of the housekeeping genes (ubiquitin C and glyceraldehyde 3-phosphate dehydrogenase, probe-based duplex primer mix, PrimerDesign) and fold change in gene expression relative to controls was determined using the ΔΔCt method. To detect SARS-CoV-2 in BCi-NS1.1, Taqman gene expression assays were used against 2019-nCoV_N1 (primers sequences from Public Health Service Centers for Disease Control and Prevention (CDC), 20 January 2020 copy) normalised to the genes HPRT, 18S and RNAse P expressions using the dCt method.

qPCR primer sequences: For amplifying long *ACE2* transcript only: Forward 5’-3’ CAAGAGCAAACGGTTGAACAC, Reverse 5’-3’ CCAGAGCCTCTCATTGTAGTCT (from Harvard PCR primer bank). For amplifying short *ACE2* transcript only: forward 5’-3’ GTGAGAGCCTTAGGTTGGATTC, reverse 5’-3’ TAAGGATCCTCCCTCCTTTGT. For amplifying both transcripts: forward 5’-3’ TGGGACTCTGCCATTTACTTAC, reverse 5’-3’ CCCAACTATCTCTCGCTTCATC

### Electrophoresis gels

All gels were 1.5% agarose in 1x TAE buffer and staining with ethidium bromide (Sigma, 5 μL per 50 ml gel). Gels were run for 75 minutes at 90 V. PCR products were loaded with 6X purple gel loading dye (B7025, NEB) and electrophoresed alongside a low molecular weight ladder (range 25 to 766 base pairs, N3233, NEB) or HyperLadder 1 kb (range 200 bp to 10 kb, BIO-33053, Bioline) to determine product sizes.

### Cilia extraction

Cilia were extracted from differentiated ALI cultures on 24-Transwell inserts (Costar) following a protocol modified from^74^. Cells on ice were washed with ice cold PBS, then incubated on the apical surface for 15 minutes in 100 μL washes with deciliation buffer (20 mM Tris hydrochloride (pH 7.5), 0.05 M sodium chloride, 10 mM calcium chloride, 1 mM EDTA, 7 mM 2-mercaptoethanol and 0.1% triton X-100)^75^ containing additional protease inhibitor cocktail (Sigma) 10uL/ml buffer. Washes were pooled and centrifuged for 2 min at 1000xg to pellet. Supernatant fractions were centrifuged at 16,000xg for 8 mins. Cilia pellets were frozen before Western blot procedue. Immunofluorescence labelling confirmed cilia enrichment, and detachment of cilia on ALI membranes. Briefly, ice cold methanol fixed and 4% dried milk blocked cilia pellets and deciliated cell membranes (excised from Transwell inserts) were labelled for 1 hour at room temperature with a mouse anti-alpha-tubulin antibody (T9026, Sigma) diluted 1:500 in PBST. Following three PBST washes a secondary goat antimouse Alexa488 antibody (Molecular Probes) was incubated for 30 minutes at room temperature before PBST washes. Deciliated cells on membranes were additionally DAPI stained before mounting. Cilia pellets and membranes were mounted in Mowiol between two glass coverslips and imaged using a Leica SP8 laser scanning confocal microscope and LAS X software.

### Western blot

Cells were washed in phosphate-buffered saline (PBS) or HBSS and lysed in 20 mM Tris-HCl pH 8.0, 137 mM NaCl, 1% (w/v) NP-40, 2mM EDTA supplemented with cOmplete™ protease inhibitor (Sigma). Samples were diluted with 5x reducing sample buffer (312.5mM Tris-HCl pH6.8, 50% glycerol, 25% 2-mercaptoethanol, 10% SDS, 0.01% bromophenol blue), incubated for 5 min at 95°C, separated on an SDS-PAGE gel and transferred to polyvinylidene fluoride membranes (BioRad). After blocking in 5% milk/TBST membranes were probed with primary anti-ACE2 antibody (Abcam 15348 for total ACE2 or Abcam 108252. for long ACE2) followed by the appropriate secondary HRP-conjugated antibody (Dako). Bound antibody was detected using Clarity ECL Western Blotting Substrate (Bio-Rad) with the image digitally captured using an Amersham Imager 600 (GE Healthcare Life Sciences). HRP-conjugated anti-β-actin antibody (Sigma) was used as a loading control. ImageJ was used for densitometry.

### Immunofluorescence cell staining

After apical wash with HBSS, cells were fixed with 4% PFA, permeabilized with 0.1% Triton X-100 and blocked with 1% BSA in PBS. Membranes were cut from the inserts and epithelial cells were stained with anti-ACE2 antibodies (ab15348 (Abcam), AF933 (R&D systems) and #NBP2-67692 (Novus)), anti-alpha tubulin (T9026 Sigma), and appropriate fluorescently labelled secondary antibodies (Alexa-488 labelled anti-mouse (Invitrogen), Alexa649 labelled anti-goat (Abcam), DyLight 647 labelled anti-rabbit (Biolegend)). Actin filaments were stained using Phalloidin-Alexafluor 555 (Cytoskeleton Inc) and nuclei with DAPI. Confocal images were taken using a Leica SP8 laser scanning confocal microscope with LAS X software.

### Antibodies

| Antigen | Species | Immunogen | Manufacturer and product code |
| --- | --- | --- | --- |
| ACE-2 | goat<br>polyclonal | Gln18-Ser740 | R&D AF933 (lot HOK0320041) |
| ACE-2 | rabbit<br>polyclonal | C-terminal domain<br>aa 788-805 | Abcam ab15348 (lot GR3333640-/-) |
| ACE-2 | rabbit<br>polyclonal | N-terminal domain<br>aa 200-300 | Abcam ab108252 (GR3338009-3) |
| Beta<br>actin | mouse<br>monoclonal | anti-β-actin-<br>peroxidase | clone AC-15, Sigma, A3854-200UL |
| Alpha tubulin | mouse monoclonal | Alpha tubulin | Sigma T9026 |

### Human rhinovirus (RV) 16 propagation and titration

Human rhinovirus (HRV16; ATCC VR-283™, Teddington, UK) was amplified using H1 HeLa cells as previously described^76,77^. Infectivity of stocks and release of infective virions in cell culture supernatants was determined using a HeLa titration assay and 50% tissue culture infective dose assay (TCID_50_/ml). Ultraviolet-irradiated virus controls (UV-RV16) were prepared by exposure of virus stocks to UV light at 1200 mJ/cm^2^ on ice for 50 min.

### Rhinovirus infection of differentiated human nasal and bronchial epithelial cells

Fully differentiated nasal and bronchial epithelial cells (28 or 21 days after ALI) were apically infected with human rhinovirus 16 (RV16) at a multiplicity of Infection (MOI) of 1 for 6h, washed apically 3x using HBSS and incubated for additional 18h at the air-liquid interface (24h in total). Cells were washed 3x with HBSS and lysed using TriZol (Invitrogen) for RNA and protein extraction.

### Interferon-treatment

PBECs monolayer cultures were stimulated with 100IU or 1000IU/ml with IFN beta (a gift from Synairgen Research Ltd) at a confluency of 70% and differentiated PBEC cultures at ALI were stimulated basolateral with 1000IU/ml IFN beta. After 24h RNA was isolated using Monarch Total RNA miniprep Kit (NEB) and reverse transcribed to cDNA using a Precision Reverse Transcription kit (PrimerDesign, Southampton, UK) according to the manufacturer’s instructions.

### Structural analysis

PyMOL Molecular Graphics System, Version 2.0 (Schrödinger, LLC) or UCSF Chimera^78^ was used to model the novel protein isoform based on full-length isoform 6M18. Molecular dynamic simulation was performed by preparing an initial 3D model of the novel protein isoform based on PDB entry 6M18 using the homology modelling function of YASARA^79^. Molecular dynamics simulations in explicit solvent were performed using YASARA with GPU acceleration^80^ on an Intel i9-9940X CPU (using 28 Threads) and GeForce RTX 2080 Ti. The molecular trajectory was sampled for 320 ns under NPT conditions at 310 K in 0.1% NaCl solution at pH 7.4 using periodic boundary conditions. Pymol, Chimera^78^ and VMD^81^ were used for molecular display and animation.

### Statistics

Statistical analyses were performed in GraphPad Prism v7.02 (GraphPad Software Inc., San Diego, CA, USA) unless otherwise indicated. For each experiment, sample size reflects the number of independent biological replicates and is provided in the figure legend. Statistical analyses of single comparisons of two groups utilized Student’s t-test or Wilcoxon Signed Rank test for parametric and non-parametric data respectively. Results were considered significant if P<0.05, where * P<0.05, ** P<0.01, *** P<0.001, ****P<0.0001.

## Contributions

VM and JSL conceptualised and supervised the initial study to interrogate RNAseq data for *ACE2* isoforms. GW identified the novel transcript, in collaboration with DED and CB, from which point CB, DED, JSL, GW and VM conceptualised and supervised the remainder of the study. CLJ, CB, GW, DED and VM designed the experiments and analysed data. CLJ and CB performed most of the experimental work with additional contributions from GW, CMS, LN, JL, FC, JC, JT, DJ, CMcC, RAR, LSND, PJS and ML. GW, CB, JSL, VM, DB, DED, RD J-MP-C, AA, PS and KT provided samples and/or resources. MF and MC developed predictive structural models. GW, VM and DED wrote the manuscript text with CB, CLJ and JSL. GW and VM prepared figures with CB, CLJ and DED. All authors approved the final submission.

## Acknowledgments

Research was supported by NIHR Southampton Biomedical Research Centre (BRC), NIHR Wellcome Trust Clinical Research Facility and AAIR Charity. GW is supported by a Wellcome Trust Seed Award in Science (204378/Z/16/Z). DB is supported by NIHR Research Professorship RP-2016-07-011. JSL, CLJ, VM, JT and JC are supported by the NHS England PCD National Service.

CB is a University of Southampton Career Track Fellow and FC is a Medical Research Foundation Fellow. ML is a BBSRC Future Leader Fellow and an NIHR Southampton BRC Senior Research Fellow.

We are grateful to healthy volunteers and respiratory patients who donated airway cells and to Synairgen Research Ltd who provided cells and reagents to support these studies.

## Data and materials availability

All sequence files to be deposited on Sequence Read Archive. Accession numbers will be provided upon publication.

---

**Supplementary Figure 1**

Boxplot and whisker showing median, quartiles and range of gene expression data for 6 selected cilia genes in primary nasal brushings (blue) and primary nasal epithelial cells cultured at ALI for 1, 4, 8, 14, 21, 28 and 63 days (orange), to demonstrate early activation of genes associated with ciliogenesis and cilium function from day 4.

**Supplementary Figure 2**

Plots of CT value against cDNA input into RT-qPCR reactions to determine dynamic range of RT-qPCRs for 3 sets of primers targeting short ACE2 only (bottom), long ACE2 (middle) and both transcripts (top).

**Supplementary Figure 3**

IF confocal images of ALI differentiated primary bronchial epithelial cells stained with anti-alpha tubulin (green), Acti-Stain 555 Phalloidin (grey), DAPI (blue) and ACE2 (red) detected with C-terminal domain antibody (top panels) or antibody detecting epitopes across the protein (middle and bottom panels), including images of red and green channels separated and merged. One higher magnification image of cells stained with the pan-ACE2 antibody is also shown (bottom)

**Supplementary Figure 4**

4a. Long ACE2 homodimer (left), long and short ACE2 heterodimer (middle) and short ACE2 homodimer (teal, from pdb 6M18) with neck dimerization domains shown in green. SARS-Cov-2 spike protein is shown in orange.

4b. DSSP analysis of short ACE2 chain A and B over the course of the 300 ns MD simulation.

4c. Analysis of helical content (left) and secondary structure variation (right) of short ACE2 over the course of the 300 ns MD simulation.

## References

1. Tipnis, S.R. et al. A human homolog of angiotensin-converting enzyme. Cloning and functional expression as a captopril-insensitive carboxypeptidase. J Biol Chem 275, 33238–43 (2000).

2. Donoghue, M. et al. A novel angiotensin-converting enzyme-related carboxypeptidase (ACE2) converts angiotensin I to angiotensin 1-9. Circ Res 87, E1–9 (2000).

3. Zhang, H. et al. Collectrin, a collecting duct-specific transmembrane glycoprotein, is a novel homolog of ACE2 and is developmentally regulated in embryonic kidneys. J Biol Chem 276, 17132–9 (2001).

4. Shulla, A. et al. A transmembrane serine protease is linked to the severe acute respiratory syndrome coronavirus receptor and activates virus entry. J Virol 85, 873–82 (2011).

5. Heurich, A. et al. TMPRSS2 and ADAM17 cleave ACE2 differentially and only proteolysis by TMPRSS2 augments entry driven by the severe acute respiratory syndrome coronavirus spike protein. J Virol 88, 1293–307 (2014).

6. Camargo, S.M. et al. Tissue-specific amino acid transporter partners ACE2 and collectrin differentially interact with hartnup mutations. Gastroenterology 136, 872–82 (2009).

7. Kowalczuk, S. et al. A protein complex in the brush-border membrane explains a Hartnup disorder allele. Faseb j 22, 2880–7 (2008).

8. Niu, M.-J., Yang, J.-K., Lin, S.-S., Ji, X.-J. & Guo, L.-M. Loss of angiotensin-converting enzyme 2 leads to impaired glucose homeostasis in mice. Endocrine 34, 56–61 (2008).

9. Bindom, S.M., Hans, C.P., Xia, H., Boulares, A.H. & Lazartigues, E. Angiotensin I-converting enzyme type 2 (ACE2) gene therapy improves glycemic control in diabetic mice. Diabetes 59, 2540–8 (2010).

10. Imai, Y. et al. Angiotensin-converting enzyme 2 protects from severe acute lung failure. Nature 436, 112–116 (2005).

11. Treml, B. et al. Recombinant angiotensin-converting enzyme 2 improves pulmonary blood flow and oxygenation in lipopolysaccharide-induced lung injury in piglets. Crit Care Med 38, 596–601 (2010).

12. Ferreira, A.J. et al. Evidence for angiotensin-converting enzyme 2 as a therapeutic target for the prevention of pulmonary hypertension. American journal of respiratory and critical care medicine 179, 1048–1054 (2009).

13. Yamazato, Y. et al. Prevention of pulmonary hypertension by Angiotensin-converting enzyme 2 gene transfer. Hypertension 54, 365–71 (2009).

14. Li, W. et al. Angiotensin-converting enzyme 2 is a functional receptor for the SARS coronavirus. Nature 426, 450–454 (2003).

15. Matsuyama, S. et al. Efficient Activation of the Severe Acute Respiratory Syndrome Coronavirus Spike Protein by the Transmembrane Protease TMPRSS2. Journal of Virology 84, 12658 (2010).

16. Kuba, K. et al. A crucial role of angiotensin converting enzyme 2 (ACE2) in SARS coronavirus–induced lung injury. Nature Medicine 11, 875–879 (2005).

17. Yan, R. et al. Structural basis for the recognition of SARS-CoV-2 by full-length human ACE2. Science (New York, N.Y.) 367, 1444–1448 (2020).

18. Wrapp, D. et al. Cryo-EM structure of the 2019-nCoV spike in the prefusion conformation. Science 367, 1260–1263 (2020).

19. Hoffmann, M. et al. SARS-CoV-2 Cell Entry Depends on ACE2 and TMPRSS2 and Is Blocked by a Clinically Proven Protease Inhibitor. Cell 181, 271–280.e8 (2020).

20. Blau, D.M. & Holmes, K.V. Human coronavirus HCoV-229E enters susceptible cells via the endocytic pathway. Adv Exp Med Biol 494, 193–8 (2001).

21. Inoue, Y. et al. Clathrin-dependent entry of severe acute respiratory syndrome coronavirus into target cells expressing ACE2 with the cytoplasmic tail deleted. Journal of virology 81, 8722–8729 (2007).

22. Wang, H. et al. SARS coronavirus entry into host cells through a novel clathrin- and caveolae-independent endocytic pathway. Cell Research 18, 290–301 (2008).

23. Pedersen, K.B., Chhabra, K.H., Nguyen, V.K., Xia, H. & Lazartigues, E. The transcription factor HNF1α induces expression of angiotensin-converting enzyme 2 (ACE2) in pancreatic islets from evolutionarily conserved promoter motifs. Biochimica et Biophysica Acta (BBA) - Gene Regulatory Mechanisms 1829, 1225–1235 (2013).

24. Kuan, T.C. et al. Identifying the regulatory element for human angiotensin-converting enzyme 2 (ACE2) expression in human cardiofibroblasts. Peptides 32, 1832–9 (2011).

25. Wang, Y. et al. Administration of 17β-estradiol to ovariectomized obese female mice reverses obesity-hypertension through an ACE2-dependent mechanism. Am J Physiol Endocrinol Metab 308, E1066–75 (2015).

26. Ziegler, C.G.K. et al. SARS-CoV-2 Receptor ACE2 Is an Interferon-Stimulated Gene in Human Airway Epithelial Cells and Is Detected in Specific Cell Subsets across Tissues. Cell 181, 1016–1035.e19 (2020).

27. The Genotype-Tissue Expression (GTEx) project. Nat Genet 45, 580–5 (2013).

28. Sungnak, W. et al. SARS-CoV-2 entry factors are highly expressed in nasal epithelial cells together with innate immune genes. Nat Med 26, 681–687 (2020).

29. Hou, Y.J. et al. SARS-CoV-2 Reverse Genetics Reveals a Variable Infection Gradient in the Respiratory Tract. Cell (2020).

30. Sims, A.C. et al. Severe acute respiratory syndrome coronavirus infection of human ciliated airway epithelia: role of ciliated cells in viral spread in the conducting airways of the lungs. Journal of virology 79, 15511–15524 (2005).

31. Dobin, A. et al. STAR: ultrafast universal RNA-seq aligner. Bioinformatics 29, 15–21 (2013).

32. Robinson, J.T. et al. Integrative genomics viewer. Nature Biotechnology 29, 24 (2011).

33. Cummings, B.B. et al. Improving genetic diagnosis in Mendelian disease with transcriptome sequencing. Sci Transl Med 9 (2017).

34. Fu, X.Y., Kessler, D.S., Veals, S.A., Levy, D.E. & Darnell, J.E., Jr. ISGF3, the transcriptional activator induced by interferon alpha, consists of multiple interacting polypeptide chains. Proceedings of the National Academy of Sciences of the United States of America 87, 8555–8559 (1990).

35. Isern, E. et al. The activator protein 1 binding motifs within the human cytomegalovirus major immediate-early enhancer are functionally redundant and act in a cooperative manner with the NF-{kappa}B sites during acute infection. J Virol 85, 1732–46 (2011).

36. Wan, F. & Lenardo, M.J. Specification of DNA binding activity of NF-kappaB proteins. Cold Spring Harbor perspectives in biology 1, a000067–a000067 (2009).

37. Walters, M.S. et al. Generation of a human airway epithelium derived basal cell line with multipotent differentiation capacity. Respiratory research 14, 135–135 (2013).

38. Jia, H.P. et al. ACE2 receptor expression and severe acute respiratory syndrome coronavirus infection depend on differentiation of human airway epithelia. Journal of virology 79, 14614–14621 (2005).

39. Wen, B., Wang, X. & Zhang, B. PepQuery enables fast, accurate, and convenient proteomic validation of novel genomic alterations. Genome Res 29, 485–493 (2019).

40. Lee, I.T. et al. Robust ACE2 protein expression localizes to the motile cilia of the respiratory tract epithelia and is not increased by ACE inhibitors or angiotensin receptor blockers. medRxiv (2020).

41. Kindler, E., Thiel, V. & Weber, F. Interaction of SARS and MERS Coronaviruses with the Antiviral Interferon Response. Adv Virus Res 96, 219–243 (2016).

42. Yuen, C.K. et al. SARS-CoV-2 nsp13, nsp14, nsp15 and orf6 function as potent interferon antagonists. Emerg Microbes Infect 9, 1418–1428 (2020).

43. Garcia-Pachon, E. et al. Asthma prevalence in patients with SARS-CoV-2 infection detected by RT-PCR not requiring hospitalization. Respir Med 171, 106084 (2020).

44. Grandbastien, M. et al. SARS-CoV-2 pneumonia in hospitalized asthmatic patients did not induce severe exacerbation. J Allergy Clin Immunol Pract (2020).

45. Chhiba, K.D. et al. Prevalence and characterization of asthma in hospitalized and non-hospitalized patients with COVID-19. J Allergy Clin Immunol (2020).

46. Camiolo, M.J., Gauthier, M., Kaminski, N., Ray, A. & Wenzel, S.E. Expression of SARS-CoV-2 Receptor ACE2 and Coincident Host Response Signature Varies by Asthma Inflammatory Phenotype. J Allergy Clin Immunol (2020).

47. Takeyama, K., Fahy, J.V. & Nadel, J.A. Relationship of epidermal growth factor receptors to goblet cell production in human bronchi. Am J Respir Crit Care Med 163, 511–6 (2001).

48. Kimura, H. et al. Type 2 inflammation modulates ACE2 and TMPRSS2 in airway epithelial cells. J Allergy Clin Immunol 146, 80–88.e8 (2020).

49. Horby, P. et al. Dexamethasone in Hospitalized Patients with Covid-19 - Preliminary Report. N Engl J Med (2020).

50. Cakebread, J.A. et al. Rhinovirus-16 induced release of IP-10 and IL-8 is augmented by Th2 cytokines in a pediatric bronchial epithelial cell model. PLoS One 9, e94010 (2014).

51. Campbell-Harding, G. et al. The innate antiviral response upregulates IL-13 receptor α2 in bronchial fibroblasts. J Allergy Clin Immunol 131, 849–55 (2013).

52. Blume, C. et al. Barrier responses of human bronchial epithelial cells to grass pollen exposure. Eur Respir J 42, 87–97 (2013).

53. Bedke, N. et al. Transforming growth factor-beta promotes rhinovirus replication in bronchial epithelial cells by suppressing the innate immune response. PLoS One 7, e44580 (2012).

54. Contoli, M. et al. Role of deficient type III interferon-lambda production in asthma exacerbations. Nat Med 12, 1023–6 (2006).

55. Wark, P.A. et al. Asthmatic bronchial epithelial cells have a deficient innate immune response to infection with rhinovirus. J Exp Med 201, 937–47 (2005).

56. Howard, T.E., Shai, S.Y., Langford, K.G., Martin, B.M. & Bernstein, K.E. Transcription of testicular angiotensin-converting enzyme (ACE) is initiated within the 12th intron of the somatic ACE gene. Mol Cell Biol 10, 4294–302 (1990).

57. Ojaghi, M., Kastelic, J. & Thundathil, J. Testis-specific isoform of angiotensin-converting enzyme (tACE) is involved in the regulation of bovine sperm capacitation. Mol Reprod Dev 84, 376–388 (2017).

58. Ho, M. Perspectives on the development of neutralizing antibodies against SARS-CoV-2. Antib Ther 3, 109–114 (2020).

59. Inal, J.M. Decoy ACE2-expressing extracellular vesicles that competitively bind SARS-CoV-2 as a possible COVID-19 therapy. Clin Sci (Lond) 134, 1301–1304 (2020).

60. Saheb Sharif-Askari, N. et al. Airways Expression of SARS-CoV-2 Receptor, ACE2, and TMPRSS2 Is Lower in Children Than Adults and Increases with Smoking and COPD. Mol Ther Methods Clin Dev 18, 1–6 (2020).

61. Pinto, B.G.G. et al. ACE2 Expression is Increased in the Lungs of Patients with Comorbidities Associated with Severe COVID-19. J Infect Dis (2020).

62. Emilsson, V. et al. ACE2 levels are altered in comorbidities linked to severe outcome in COVID-19. medRxiv (2020).

63. Zhou, L. et al. ACE2 and TMPRSS2 are expressed on the human ocular surface, suggesting susceptibility to SARS-CoV-2 infection. Ocul Surf 18, 537–44 (2020).

64. Collin, J. et al. Co-expression of SARS-CoV-2 entry genes in the superficial adult human conjunctival, limbal and corneal epithelium suggests an additional route of entry via the ocular surface. Ocul Surf (2020).

65. Lange, C. et al. Expression of the COVID-19 receptor ACE2 in the human conjunctiva. J Med Virol (2020).

66. Ma, D. et al. Expression of SARS-CoV-2 receptor ACE2 and TMPRSS2 in human primary conjunctival and pterygium cell lines and in mouse cornea. Eye (Lond), 1–8 (2020).

67. Zhang, D. et al. Incomplete annotation has a disproportionate impact on our understanding of Mendelian and complex neurogenetic disorders. Science Advances 6, eaay8299 (2020).

68. Hirst, R.A. et al. Culture of primary ciliary dyskinesia epithelial cells at air-liquid interface can alter ciliary phenotype but remains a robust and informative diagnostic aid. PLoS One 9, e89675 (2014).

69. Xiao, C. et al. Defective epithelial barrier function in asthma. J Allergy Clin Immunol 128, 549–56.e1-12 (2011).

70. Kuek, L.E. et al. Identification of an Immortalized Human Airway Epithelial Cell Line with Dyskinetic Cilia. Am J Respir Cell Mol Biol 59, 375–382 (2018).

71. Wang, L., Wang, S. & Li, W. RSeQC: quality control of RNA-seq experiments. Bioinformatics 28, 2184–5 (2012).

72. Shao, M. & Kingsford, C. Accurate assembly of transcripts through phase-preserving graph decomposition. Nature Biotechnology 35, 1167–1169 (2017).

73. Patro, R., Duggal, G., Love, M.I., Irizarry, R.A. & Kingsford, C. Salmon provides fast and bias-aware quantification of transcript expression. Nat Methods 14, 417–419 (2017).

74. Ostrowski, L.E. et al. A Proteomic Analysis of Human Cilia. Molecular & Cellular Proteomics 1, 451 (2002).

75. Hastie, A.T. et al. Isolation of cilia from porcine tracheal epithelium and extraction of dynein arms. Cell Motil Cytoskeleton 6, 25–34 (1986).

76. Calvén, J. et al. Viral stimuli trigger exaggerated thymic stromal lymphopoietin expression by chronic obstructive pulmonary disease epithelium: role of endosomal TLR3 and cytosolic RIG-I-like helicases. J Innate Immun 4, 86–99 (2012).

77. Zhao, W. et al. Peroxisome proliferator-activated receptor gamma negatively regulates IFN-beta production in Toll-like receptor (TLR) 3- and TLR4-stimulated macrophages by preventing interferon regulatory factor 3 binding to the IFN-beta promoter. J Biol Chem 286, 5519–28 (2011).

78. Pettersen, E.F. et al. UCSF Chimera--a visualization system for exploratory research and analysis. J Comput Chem 25, 1605–12 (2004).

79. Krieger, E. et al. Improving physical realism, stereochemistry, and side-chain accuracy in homology modeling: Four approaches that performed well in CASP8. Proteins 77 Suppl 9, 114–22 (2009).

80. Krieger, E. & Vriend, G. New ways to boost molecular dynamics simulations. J Comput Chem 36, 996–1007 (2015).

81. Humphrey, W., Dalke, A. & Schulten, K. VMD: visual molecular dynamics. J Mol Graph 14, 33–8, 27–8 (1996).

