## Supplementary figures and images for "A novel isoform of *ACE2* is expressed in human nasal and bronchial respiratory epithelia and is upregulated in response to RNA respiratory virus infection"

### Supplementary Figure 1

#### Supplemental Figure 1

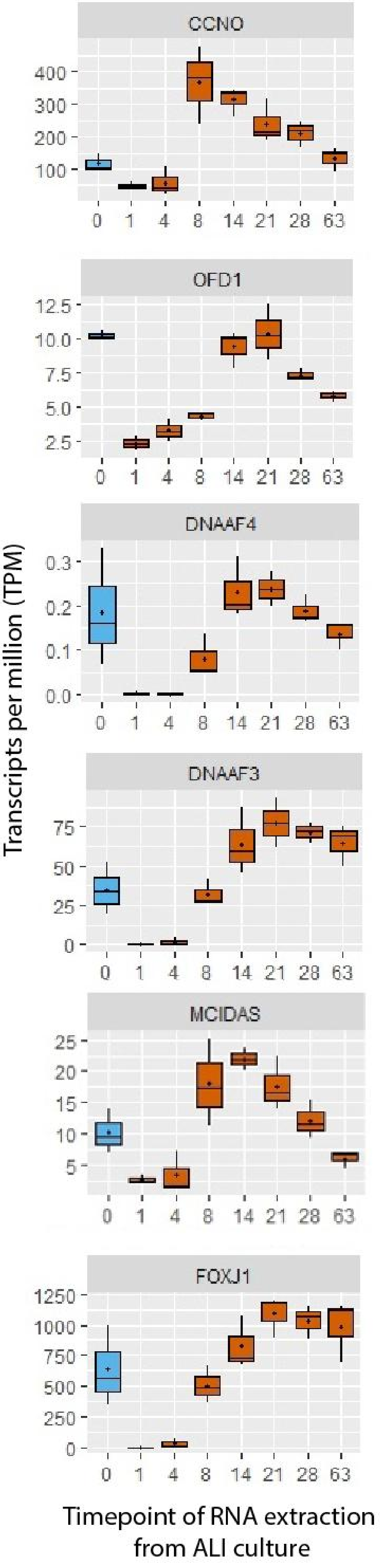

### Supplementary Figure 2

#### Supplemental Figure 2

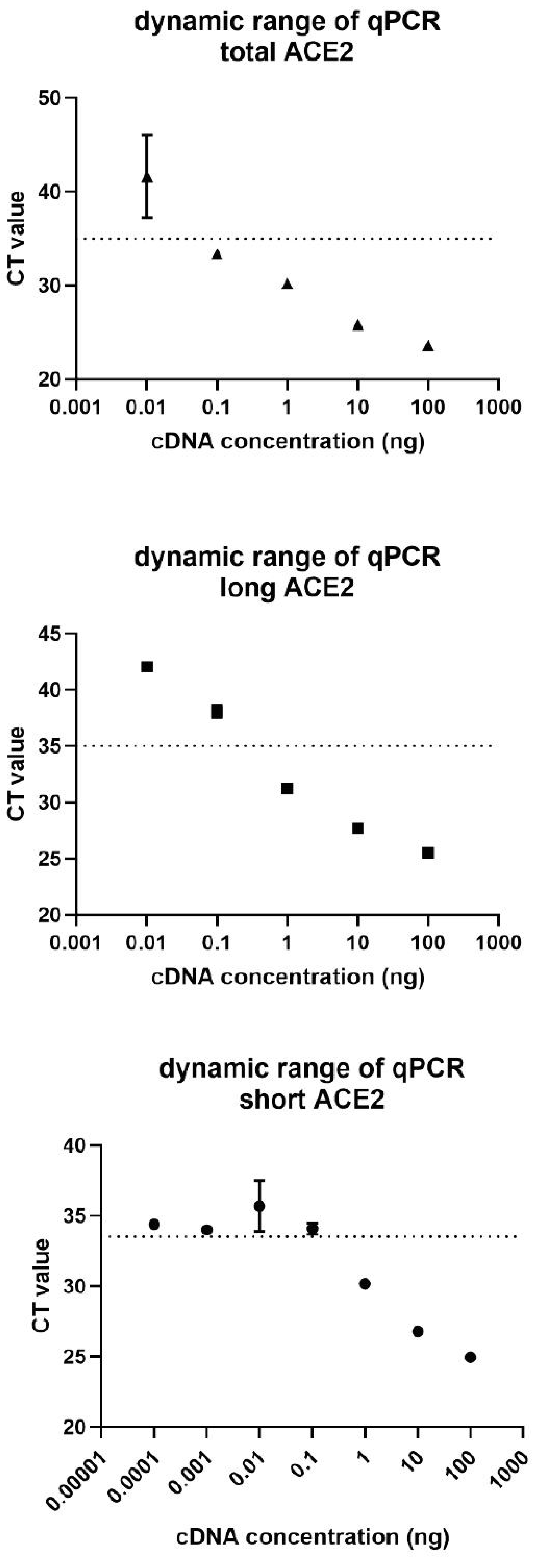

### Supplementary Figure 3

#### Supplemental Figure 3

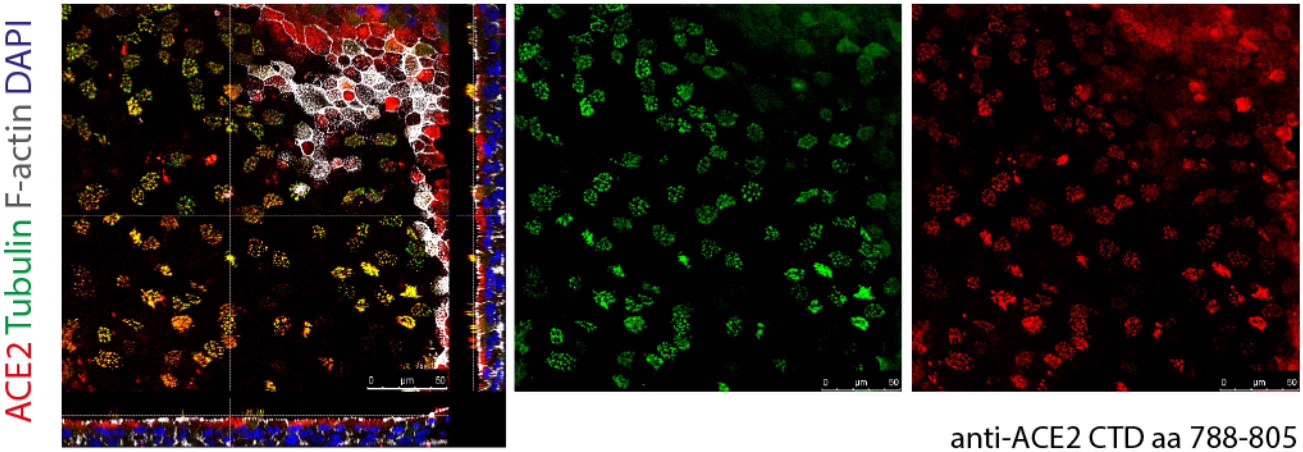


anti-ACE2 CTD aa 788-805
Abcam ab15348

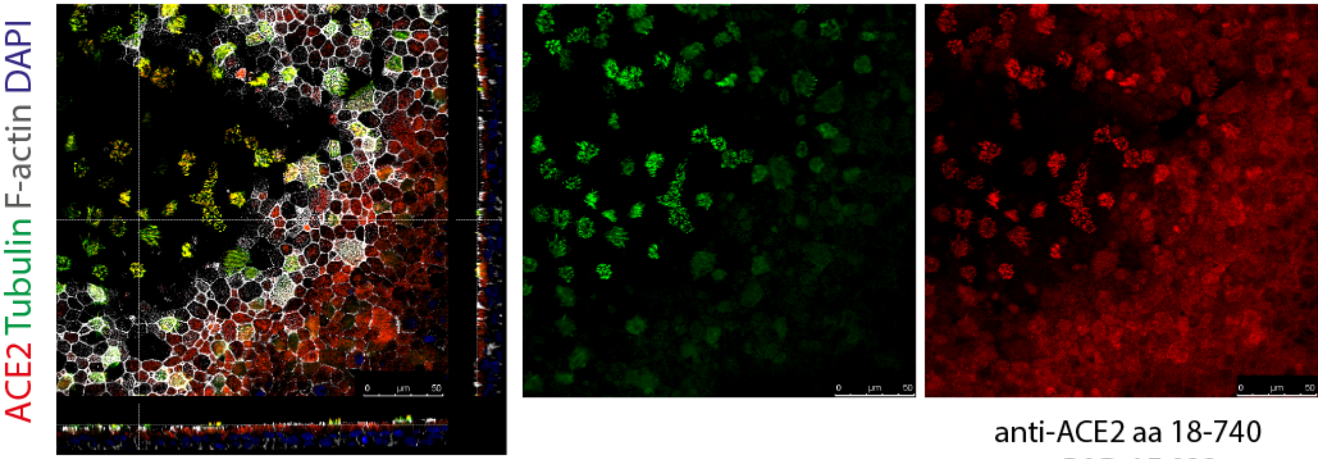


anti-ACE2 aa 18-740
R&D AF 933

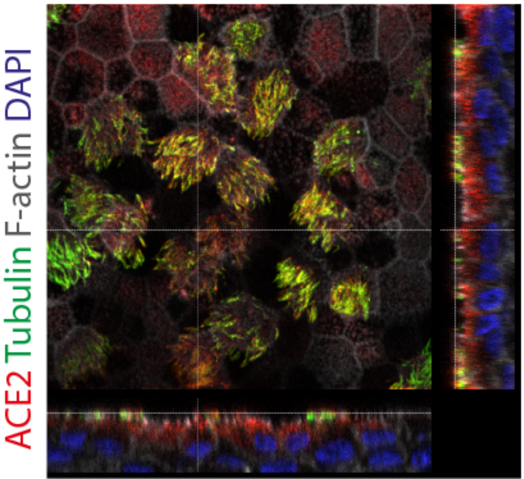


anti-ACE2 aa 18-740
R&D AF 933

### Supplementary Figure 4

#### Supplemental Figure 4

a

LONG ACE2 HOMODIMER | LONG-SHORT ACE2 HETERODIMER | SHORT ACE2 HOMODIMER

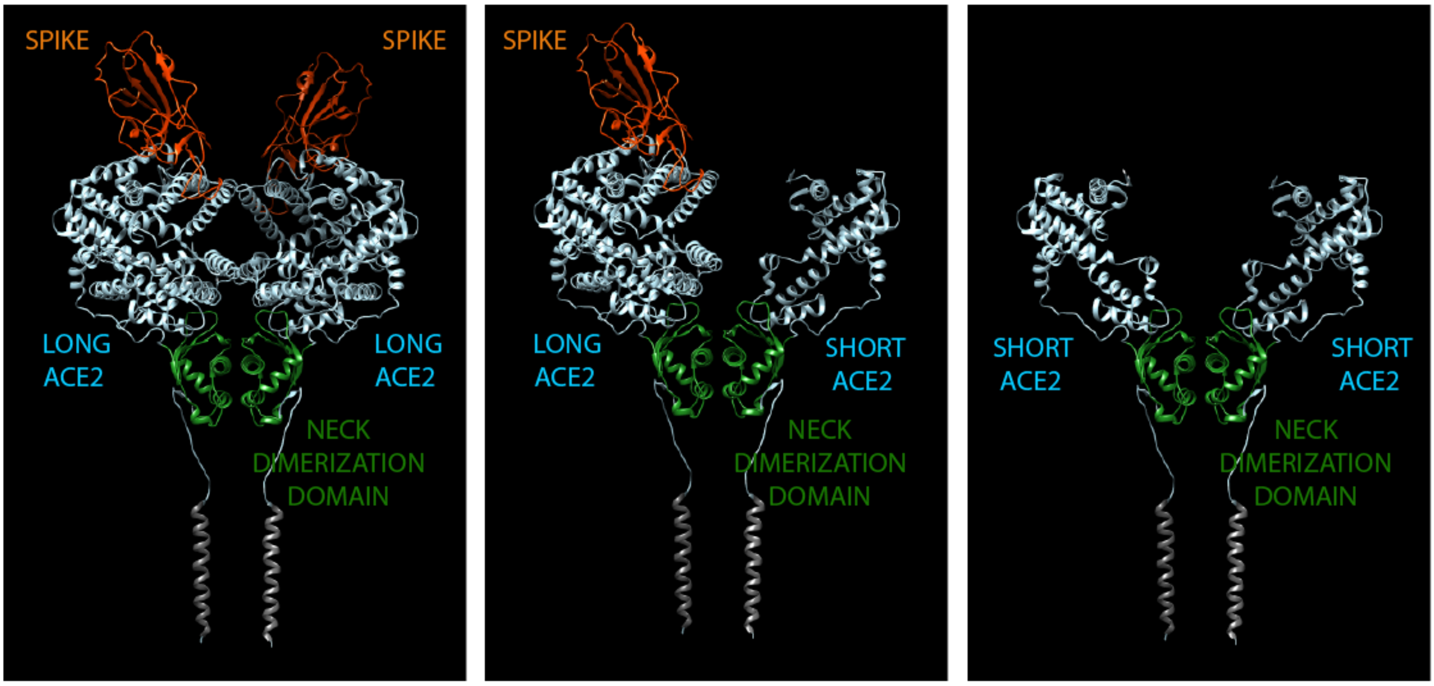


b

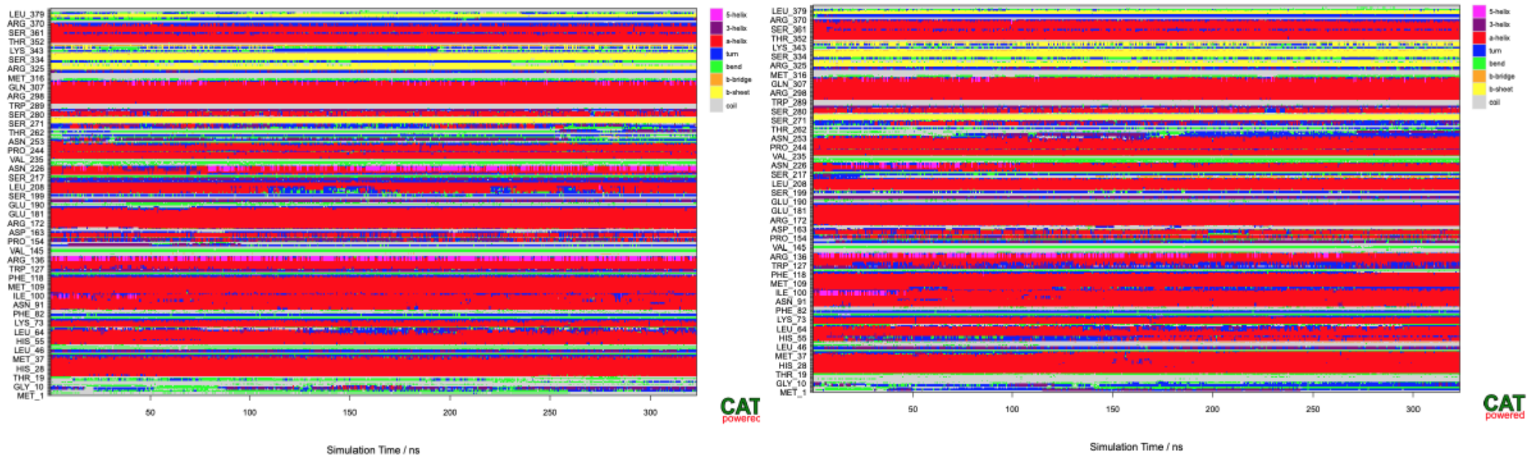


c

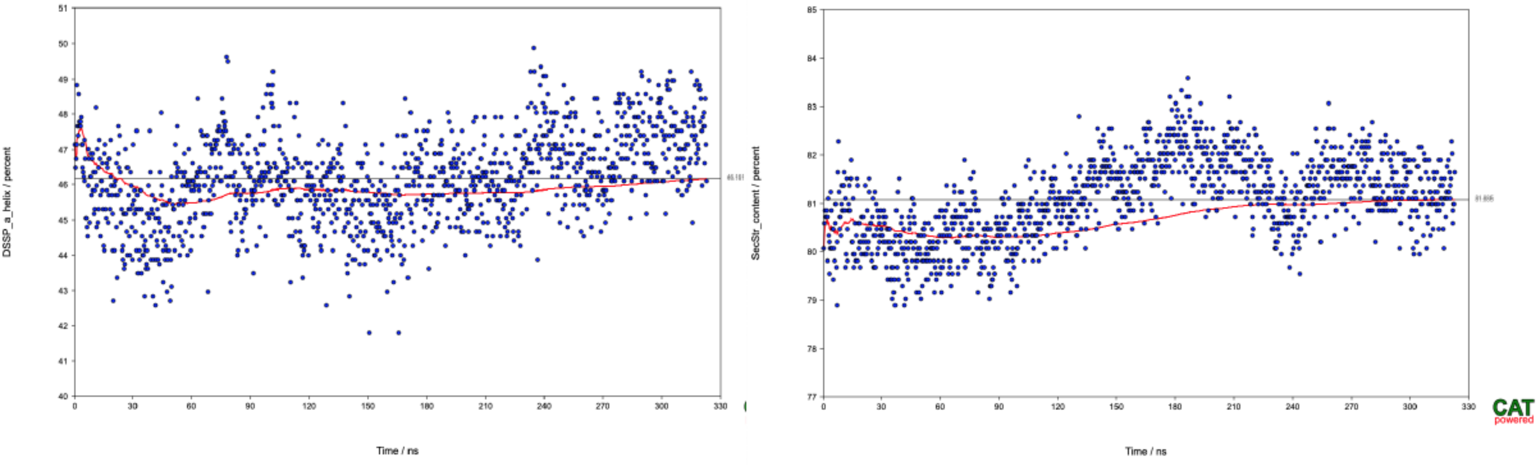
